# Differentiation of exhausted CD8 T cells after termination of chronic antigen stimulation stops short of achieving functional T cell memory

**DOI:** 10.1101/2021.04.03.437705

**Authors:** Pierre Tonnerre, David Wolski, Sonu Subudhi, Jihad Al-Jabban, Ruben C. Hoogeveen, Marcos Damasio, Hannah K. Drescher, Lea M. Bartsch, Damien C. Tully, Debattama R. Sen, David J. Bean, Joelle Brown, Almudena Torres-Cornejo, Maxwell Robidoux, Daniel Kvistad, Nadia Alatrakchi, Ang Cui, David Lieb, James A. Cheney, Jenna Gustafson, Lia L Lewis-Ximenez, Lucile Massenet-Regad, Thomas Eisenhaure, Jasneet Aneja, W. Nicholas Haining, Raymond T. Chung, Nir Hacohen, Todd M. Allen, Arthur Y. Kim, Georg M. Lauer

**Affiliations:** Division of Gastroenterology, Massachusetts General Hospital and Harvard Medical School, Boston, MA, USA; Inserm U976, Institut de Recherche Saint-Louis, Paris, France; Ragon Institute of MGH, MIT and Harvard, Cambridge, MA, USA; Division of Medical Sciences, Harvard Medical School, Boston, MA, USA; Department of Pediatric Oncology, Dana-Farber Cancer Institute, Boston, MA, USA; Broad Institute of MIT and Harvard, Cambridge, MA, USA; Harvard-MIT Division of Health Sciences and Technology, MIT, Cambridge, MA, USA; Instituto Oswaldo Cruz, Fundação Oswaldo Cruz, Rio de Janeiro, Brazil; Center for Cancer Research, Massachusetts General Hospital, Boston, MA, USA; Division of Infectious Diseases, Massachusetts General Hospital and Harvard Medical School, Boston, MA, USA

**Keywords:** T cell exhaustion, immunological recovery, HCV infection, antiviral therapy

## Abstract

T cell exhaustion is associated with failure to clear chronic infections and malignant cells. Defining the molecular mechanisms of T cell exhaustion and reinvigoration is essential to improving immunotherapeutic modalities. Analysis of antigen-specific CD8+ T cells before and after antigen removal in human hepatitis C virus (HCV) infection confirmed pervasive phenotypic, functional, and transcriptional differences between exhausted and memory CD8+ T cells. After viral cure, we observed broad phenotypic and transcriptional changes in clonally stable exhausted T-cell populations suggesting differentiation towards a memory-like profile. However, functionally, the cells showed little improvement and critical transcriptional regulators remained in the exhaustion state. Notably, T cells from chronic HCV infection that were exposed to antigen for shorter periods of time because of viral escape mutations were functionally and transcriptionally more similar to memory T cells from spontaneously resolved acute HCV infection. Thus, duration of T cell stimulation impacts the ability to recover from exhaustion, as antigen removal after long-term T cell exhaustion is insufficient for the development of key T cell memory characteristics.

## Introduction

Chronic viral infections and cancer are characterized by accumulating changes in antigen-specific CD8+ T cells, termed T cell exhaustion^1, 2^. T cell exhaustion is initiated and maintained by extended exposure to cognate antigen and inflammatory signals^2, 3^. Exhausted T cells (T_EX_) characteristically express inhibitory receptors such as PD-1 or 2B4^4–6^, increasingly lose key functions such as cytokine secretion and proliferation^3, 7^, and do not differentiate into memory T cells (T_MEM_) as after acute infection^8^. T_EX_ also lack antigen-independent self-renewal^9, 10^ and the ability to mount a swift recall response. At the extreme end of the exhaustion spectrum antigen-specific T cells even get physically deleted^9, 11^.

Checkpoint inhibitor therapies targeting PD-1 and CTLA-4 have revolutionized cancer therapy, demonstrating that T cell exhaustion can be overcome^12, 13^. However, current therapies are only effective for some cancers and, within a disease category, work only for select patient groups, indicating that the molecular mechanisms underlying T-cell exhaustion are complex and heterogeneous^14^. Additionally, immune recovery is not long lasting^15^. In addition to checkpoint inhibition, other treatment modalities aiming at invigorating T cell responses, such as therapeutic vaccines^16, 17^ as well as immunomodulatory or antigen lowering drugs in chronic viral infection^18^, will rely on the ability to overcome the dysfunctionality of exhausted T cells. Understanding the cellular pathways driving T-cell exhaustion and the road to sustained T cell recovery is critical for developing more effective and targeted therapies^14^.

While the major clinical breakthroughs based on reversing T cell exhaustion have been achieved in cancer, much of our understanding of the molecular mechanisms of T cell exhaustion stems from studies of chronic viral infection, most notably with different strains of LCMV in mice^19^. The major advantage of studying viral infection is that it readily allows analysis of T cell exhaustion in antigen-specific T cells and within the critical context of antigen burden, both of which are much more difficult to define in cancer. Emerging tools to study small populations of immune cells from clinical samples enable studies of T cell exhaustion directly in humans, especially in chronic infections with HIV^20^, hepatitis B virus (HBV)^21, 22^, and hepatitis C virus (HCV)^4, 23^. HCV infection is particularly suited to elucidating mechanisms of T cell differentiation in acute and chronic settings. HCV is the only chronic viral infection with a complete dichotomy in natural outcome^24^, as an estimated 20-30% of infected subjects completely clear the virus, typically within 6 months, allowing direct comparison of T cell memory and exhaustion within the same pathogen and host framework^25^. Additionally, HCV is the first, and so far only, chronic viral infection that can be completely terminated using direct-acting antivirals (DAA)^26^, enabling study of whether termination of antigen exposure enables exhausted T cell populations to re-differentiate into effective T cell memory. Such work has revealed important insights, including T cell proliferation recovery after antigen clearance in parallel to the expansion of pre-existing CD127+ PD-1+ TCF-1+ memory-like T cells^27, 28^. Nevertheless, these memory-like CD8+ T cells are partially different from actual T_MEM_ after acute infection^27, 29, 30^, indicating the need for more detailed analyses to understand the molecular trajectories of T_EX_ after antigen removal and the hurdles to achieving full memory potential.

In this study, we utilized the paradigm of chronic HCV infection and DAA treatment to further define the features of T cell exhaustion in humans, and the potential for exhaustion to revert after antigen removal. Through a specifically designed clinical DAA trial that incorporated leukapheresis collection at multiple timepoints we were able to perform broad and deep T cell studies using samples from the same peripheral blood mononuclear cell (PBMC) collections in the context of a very well-defined clinical perturbation. After cure, we documented pervasive changes in clonally stable exhausted T cell populations suggesting differentiation towards a more memory like phenotype, while function and critical transcriptional regulators were mostly fixed in the exhausted T cell state. We also provide evidences that this T cell scarring solidifies with the duration of exhaustion, indicating a limited window of opportunity early in chronic infection.

## Results

### An HCV DAA trial designed to test whether elimination of viral antigen enables differentiation of T_EX_ into effective CD8+ T cell memory

To determine the impact of antigen removal in a state of chronic viral infection, we designed a DAA treatment trial for HCV infection, generating optimal samples for immunological studies. We were able to study 20 of 25 long-term HCV-infected patients participating in the trial and receiving 12 weeks of paritaprevir/ritonavir/ombitasvir + dasabuvir + ribavirin^31^ (Extended Data **Table 1**), with structured PBMC collection pre- and post-therapy, including at least two leukapheresis collections at weeks 0 and 24. A typical yield of 10^10^ PBMCs per leukapheresis procedure allowed us to perform all CD8+ T cell assays reported in this manuscript on cells from the same research sample and on large virus-specific T cell populations, despite the low frequency of HCV-specific CD8+ T cells in the blood.

First, we screened all patients for the presence of HCV-specific CD8+ T cells using HLA class I multimers matching their expressed HLA alleles at baseline and 12 weeks after cessation of treatment (**Fig. 1a** and **Extended Data Fig. 1a,b**). Even extremely low-frequency HCV-specific T cell populations could be analyzed after magnetic bead enrichment of multimer-positive cells from up to 10^8^ PBMCs. After DAA treatment these low frequencies remained stable or declined to even lower levels (**Extended Data Fig. 1c,d**). In parallel, we also determined the sequence of each targeted epitope in the viral populations circulating in each patient using next generation viral sequencing (**Fig. 1b** and **Extended Data Fig. 2a**). Based on these results we subsequently tested whether the identified CD8+ T cell responses were actually able to recognize the currently dominant viral variants (**Extended Data Fig. 2b**). This is critical for T cell assay interpretation, since, similar to HIV, HCV has the ability to escape T cell recognition via the emergence of viral variants that impede T-cell receptor (TCR) binding or activation. Importantly, and in contrast to HIV, viral escape from T cells in HCV is only observed in the early phase of infection, i.e., within a year after exposure^32, 33^, meaning that T cells targeting escaped epitopes received antigen signal for only a limited time, with limited or no TCR signal afterwards, while remaining in the same inflammatory environment. This makes T cells targeting escaped epitopes suitable controls for the impact of the duration of TCR stimulation on T cells. We could detect T cell responses targeting both preserved and escaped T cell epitopes in most patients (**Fig. 1b** and **Extended Data Fig. 2**) and classified them as follows. When the epitope sequence matched the sequence of the circulating virus, we considered these T cells with full recognition of the autologous virus as likely to be exhausted and therefore labeled them T_EX_. If viral sequencing detected a dominant population of viral variants different from the prototype epitope, we performed functional testing of HCV-specific CD8+ T cells with both wild-type and variant peptides. This allowed us to determine whether viral variants completely abrogated peptide recognition or only diminished the T cell response (**Fig. 1b** and **Extended Data Fig. 2**). The complete abrogation of a response against a viral variant can only be explained by a priming with the prototype sequence followed by viral evolution, i.e. viral escape mutation, and thus these T cell responses were categorized as full escape, or T_F-ESC_^23, 32–34^. Variants with partial recognition by the T cells were labeled as partial escape, or T_P-ESC_^33^. For T_P-ESC_ sequences we cannot completely rule out that they were already primed by the variant sequence, and thus we focused most of our analyses on T_EX_ and T_F-ESC_. Importantly, different viral recognition patterns were associated with distinct expression patterns of memory- and exhaustion-related molecules, supporting the classification of T cell responses as proposed (**Fig. 1c**). Furthermore, we also observed that T_EX_ frequencies typically decreased after viral control, whereas frequencies of T cells targeting escaped epitopes remained stable or even slightly increased post-therapy (**Extended Data Fig. 1d**), further substantiating the difference in TCR signaling depending on the current epitope sequence. As non-HCV controls, we identified CD8+ T cell populations targeting different acute and chronic viruses (influenza, EBV and CMV) that also would not undergo changes in antigen stimulation levels during HCV DAA therapy (**Fig. 1b**). An example of the different virus-specific CD8+ T cells identified in one individual can be found in **Fig. 1d**.

**Figure 1:**
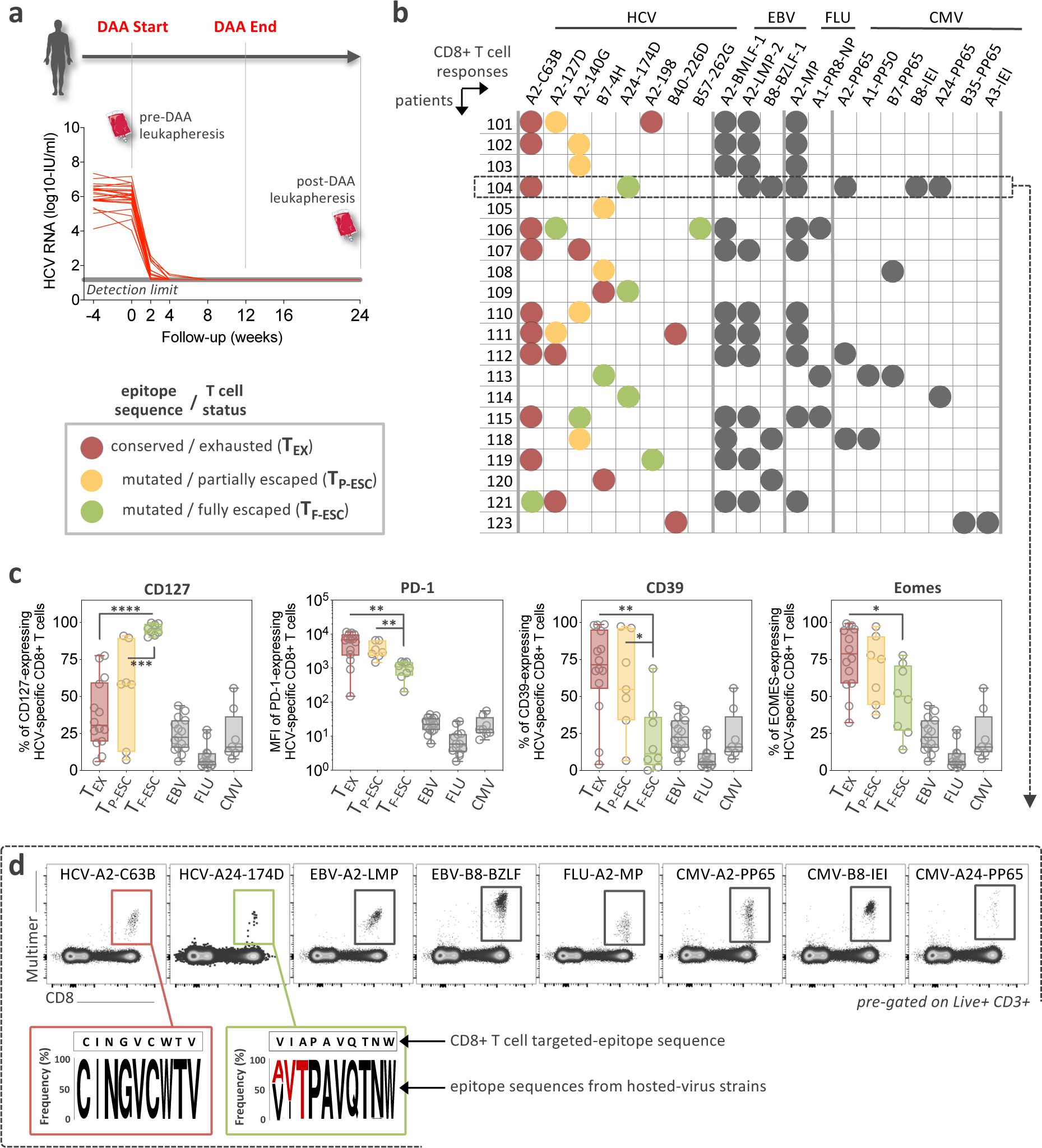
Study design, patients and virus-specific CD8+ T cell responses. **a**, Overview of the clinical trial phases and longitudinal monitoring of HCV viral load. **b,** Study-subject virus-specific CD8+ T cell responses detected by MHC class I multimers. Targeted HCV-epitopes were sequenced and grouped as “conserved / exhausted (T_EX_)” (red circles), “mutated / partially escaped (T_P-ESC_)” (orange circles), or “mutated / fully escaped (T_F-ESC_)” (green circles) according to subsequent testing of the T cell recognition of the variant compared to wild-type epitopes through intracellular staining of IFNγ (Extended Data Fig, 2). **c**, Dot plot histograms displaying CD127, PD-1, CD39 and Eomes expression by HCV-specific T_EX_, T_P-ESC_ and T_F-ESC_ as well as by EBV-, FLU- and CMV-specific T cells, pre-DAA therapy. Statistical testing by Mann-Whitney tests (unpaired, nonparametric). **d,** Representative flow cytometry plots of virus-specific CD8+ T cells from patient 104, and related HCV epitope-sequences.

**Figure 2:**
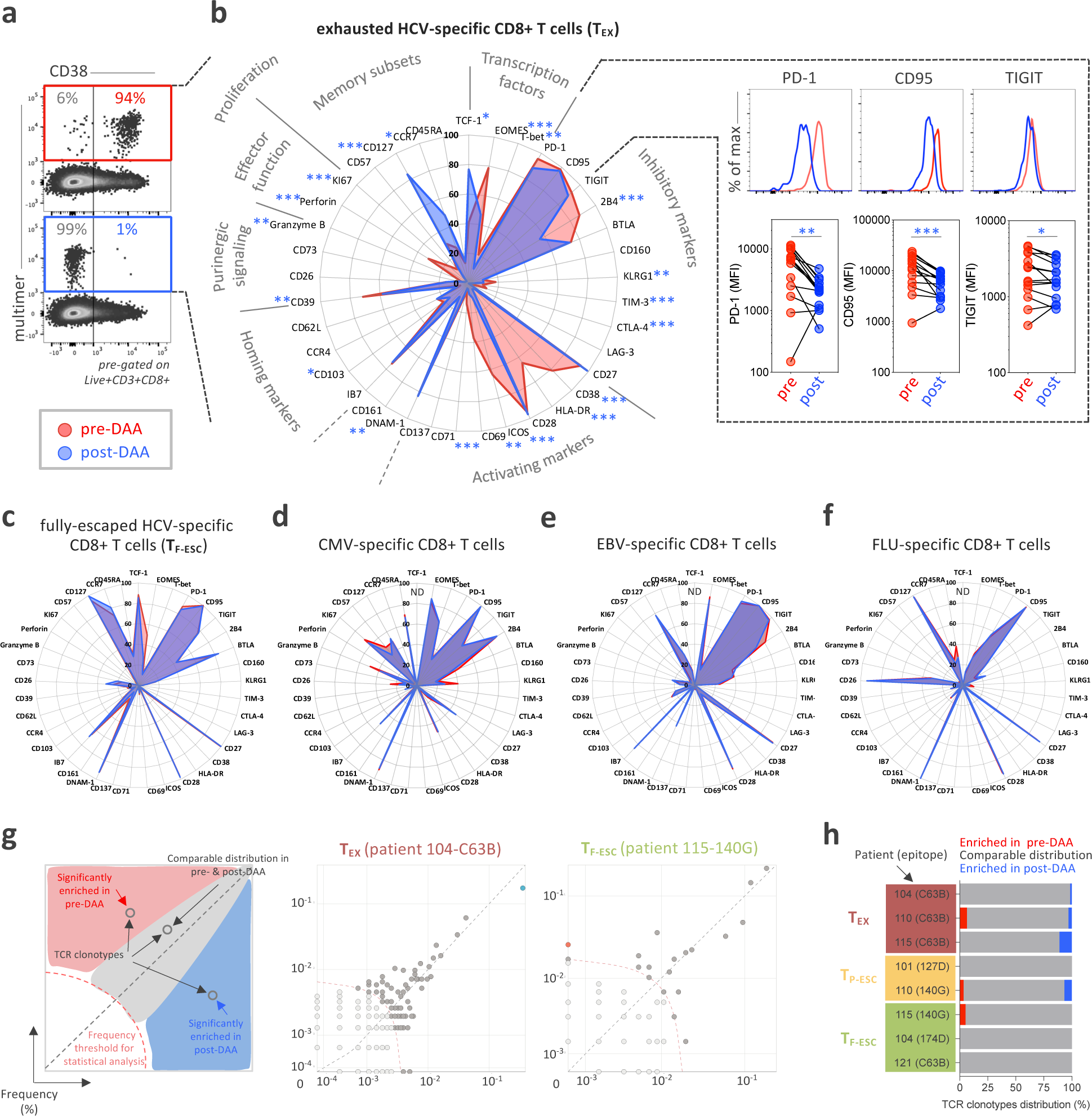
HCV-specific T_EX_ have a characteristic phenotype that is significantly changed after DAA therapy and antigen removal. **a**, Representative flow cytometry dot plots of the changes occurring in the expression of CD38 pre- and post-DAA therapy. **b**, Deep immune-profiling analysis of T_EX_ HCV-specific CD8+ T cells (paired samples, n=14) before (red plot) and after (blue plot) DAA-treatment. Data are expressed as percentage of expressing cells for the different markers listed. Changes in expression level, defined by the median fluorescence intensity (MFI) for PD1, CD95 and TIGIT before and after DAA treatment are also indicated in the right panels. Statistical testing by Wilcoxon tests (paired, nonparametric). **c**, Phenotypic changes in T_F-ESC_ HCV-specific CD8+T cells targeting mutated epitopes (paired samples, n=8) pre- and post-DAA treatment. Phenotypic changes in CMV- (paired samples, n=8), EBV- (paired samples, n=14), and influenza-specific (paired samples, n=13) CD8+T cells pre- and post-DAA treatment are displayed in **d**, **e** and **f**, respectively. **g**, Correlation plots displaying the frequencies of the different clonotypes identified by TCR sequencing between pre- and post-DAA therapy. Each dot represents a unique TCR clonotype. **h**, TCR clonotypes distribution in n=8 different T cell populations (T_EX_, n=3; T_P-ESC_, n=2; and T_F-ESC_, n=3) between pre- and post-DAA treatment.

### DAA therapy induces characteristic phenotypical changes in clonally stable HCV-specific T_EX_

Based on the CD8+ T cell response screening, we performed comprehensive immune profiling by flow cytometry pre- and post-DAA therapy, assessing the expression of 37 different molecules with known relevance for T cell state and differentiation (**Fig. 2a,b** and **Extended Data Fig. 3a-c)**. HCV-specific CD8+ T cells targeting conserved epitopes (T_EX_) had the expected profile of high activation with expression of CD38, HLA-DR, ICOS, and CD69 by most HCV-specific T cells, an effector memory phenotype (CCR7_lo_, CD45RA_lo_, and CD127_lo,_), high expression of different T cell inhibitory receptors and other molecules related to T cell exhaustion (PD-1, TIGIT, CD95, BTLA, 2B4, and CD39), and the typical transcription factor profile (TCF-1_lo_, Eomes_hi_, and T-bet_lo_) (**Fig. 2b**, red line). This exhausted T cell phenotype had changed extensively by 12 weeks after the end of DAA therapy, or almost 24 weeks after termination of viremia, with 23/37 molecules expressed at significantly different levels. The overall picture post HCV cure was characterized by a complete reduction in T cell activation (complete loss of CD38, HLA-DR, ICOS, CD69 and CD71 expression), a switch toward a central memory phenotype with more cells expressing CCR7 and, especially, CD127, and a switch towards a higher frequency of TCF-1 than Eomes-expressing cells (**Fig. 2b**, blue line). While most HCV-specific CD8 T_EX_ cells continued to express T cell inhibitory molecules, most T cell inhibitory molecules were significantly decreased in either percentage (2B4, CD39, TIM-3, and CTLA-4) or median fluorescence intensity (MFI) (PD-1, CD95, and TIGIT) (**Fig. 2b**, right panels). Overall, the phenotypic changes in T_EX_ post-DAA therapy confirmed a switch towards less-exhausted T cells that had developed distinct features usually associated with T_MEM_. All other virus-specific CD8+ T cell populations, including HCV-specific T_F-ESC_ and those targeting other viruses, displayed a very different phenotypic T cell profile at baseline, and underwent no significant changes from pre- to post-DAA therapy (**Fig. 2c-f**). There was also no change in the phenotype of bulk CD8 T cell populations (data not shown). Together, the data expand on previous findings^27^ by describing even broader changes from the exhaustion phenotype towards memory features. Further, even when studying 37 different T cell parameters, all observed changes can be attributed to the removal of HCV antigen, and thus cessation of TCR stimulation, rather than the termination of chronic inflammation. In addition, we performed TCR sequencing analyses in a subset of patients in order to determine whether the broad phenotypical changes described above could be explained by a remodeling of the clonal composition of T_EX_ post-therapy. As we observed rather stable clonal repertoires in both T_EX_ and T_ESC_ (**Fig. 2g,h**), the pervasive phenotypic changes in T_EX_ cannot be explained by selective emergence or disappearance of distinct HCV-specific T cell clones, but rather by differentiation of cells from within the original clonal repertoire.

**Figure 3:**
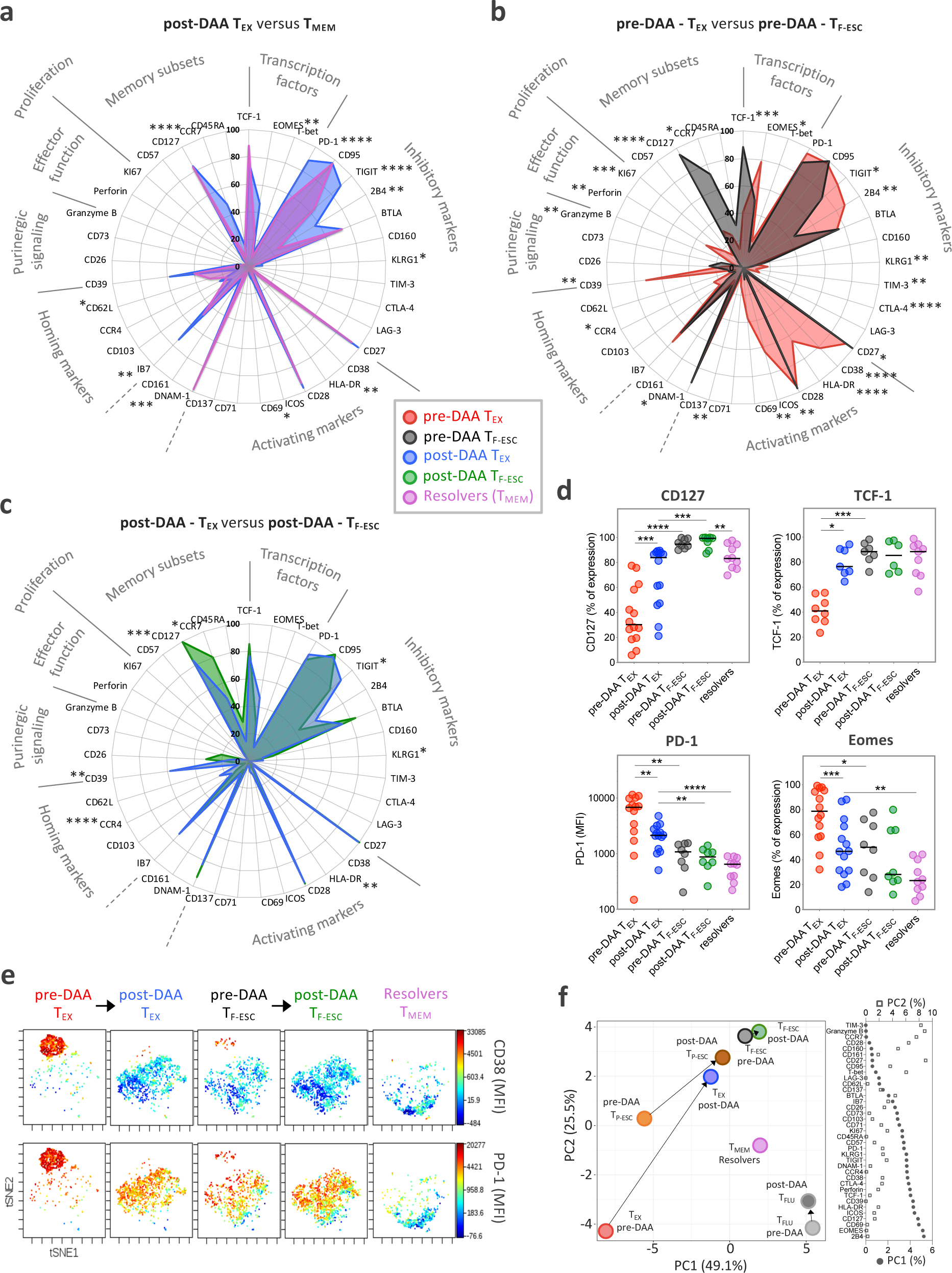
The phenotypic change of T_EX_ towards T_MEM_ after HCV cure remains incomplete. **a**, Comparison of the phenotypic immune signature of post-DAA T_EX_ (n=14) to that of T_MEM_ from spontaneous HCV-resolvers (n=10). **b**, Comparison of the T_EX_ phenotypic immune signature (n=14) to that of T_F-ESC_ (n=8) pre-DAA therapy. **c**, Comparison of the T_EX_ phenotypical immune-signature (n=14) to T_F-ESC_ (n=8), post-DAA therapy. **a-c**, Data are expressed as the percentage of cells expressing the listed markers. Dot plot histograms comparing expression levels of the 37 proteins studied across the different HCV-specific CD8+ T cell populations are presented in Extended Data Fig. 3b. **d**, Dot plot histograms displaying CD127, TCF-1, PD-1 and Eomes expression levels across T_EX_ and T_F-ESC_, pre- and post-DAA therapy, and in resolver T_MEM._ **a-d**, Comparison rules and statistical tests are as presented in Extended Data Fig.3c. **e**, t-SNE analysis of T_EX_ and T_F-ESC_, pre- and post-DAA therapy, as well as resolver T_MEM_, based on the expression levels of CD38, HLA-DR, PD-1, CD39, TIGIT, CCR7, CD45RA, Integrin-Beta-7 and CD62L. Expression levels (MFI) of CD38 and PD-1 are displayed using a color scheme. **f**, Principal component analysis of the expression levels of the 37 proteins, as detected by flow cytometry, by T_EX_, T_P-ESC_, T_F-ESC_ and T_FLU_, pre- and post-DAA therapy, as well as by resolver T_MEM_. Respective contribution of the 37 different proteins in driving PC1 and PC2 are depicted in the right panel.

### The phenotypical transformation of T_EX_ towards T_MEM_ after antigen clearance remains incomplete

We next tested whether the phenotypic changes in T_EX_ after cure of chronic infection resulted in T cell populations recapitulating true T cell memory by comparing their phenotype to HCV-specific CD8+ T cells from patients who had spontaneously resolved HCV infection (**Extended Data Fig. 1e**). In patients with a recently resolved acute infection, we analyzed PBMCs from about 24 weeks after the last positive viral load, so that the HCV-specific CD8+ T cells were at a similar time since the last TCR signal as the T_EX_ after cure. The expression levels of some molecules, such as CD127 and TCF-1, became similar to those of T_MEM_ (**Fig. 3a,d** and **Extended Data Fig. 3b,c**). Other important molecules, however, remained differentially expressed, including PD-1 and Eomes (**Fig. 3d** and **Extended Data Fig. 3b,c**). When we added CD8+ T cells targeting escape epitopes to the analysis, differences between T_EX_ and T_F-ESC_ narrowed from pre- to post-HCV cure, but significant phenotypic differences also remained between these T cell populations (**Fig. 3b-d** and **Extended Data Fig. 3b,c**). Overall, T_F-ESC_ occupied a space between T_EX_ after cure and T_MEM_ (**Fig. 3b-d** and **Extended Data Fig. 3b,c**). This finding was further supported through T-distributed stochastic neighbor embedding (t-SNE) visualization based on the expression levels of 9 different molecules that were part of one of the flow cytometry phenotyping panels (**Fig. 3e**, expression levels shown for CD38 and PD-1). Looking at individual T cells from the different HCV-specific T cell populations, we observe two highly distinct clusters at opposite sides of the spectrum. In the upper left corner, we almost exclusively see T_EX_ from pre-treatment, with high PD-1 and CD38 expression (**Fig. 3e, left panel**). In contrast, cells in the lower right corner are completely negative for CD38 and PD-1 and consist only of T_MEM_ from resolved infections (**Fig. 3e, right panel**). T_EX_ after treatment move towards the lower right, but fall significantly short of reaching the t-SNE space with a full T_MEM_ profile (second panel from left). T_ESC_ from both pre- and post-treatment (middle panel and second panel from right) also occupy a space in the middle, but reach a position closer to T_MEM_. Similar results were observed after principal component (PC) analysis of the same data used for t-SNE analysis (**Extended Data Fig. 3d**), and this pattern was further substantiated by PC analysis integrating the results from all 37 molecules that we analyzed via flow cytometry (**Fig. 3f** and **Extended Data Fig. 3e**). Pre-treatment, we see increasingly more memory-like T cells on the trajectory from T_EX_ over T_P-ESC_ to T_F-ESC_ on both the PC1 and PC2 axis, with further differentiation towards memory post cure status, though none of the T cell populations from chronic infection fully reach the memory T cell profile on the bottom right of the PC plane, where T_MEM_ from HCV-resolvers and Flu-specific T cells are located. Together, the data indicate that T_EX_ do not fully differentiate into T_MEM_ post removal of antigen, but rather acquire a memory-like phenotype similar to what we observe in T_ESC_ already after viral escape early in infection.

### T_ESC_, but not T_EX_ after viral cure, display functional properties similar to those of T_MEM_

Despite broad phenotyping, the quality of a T cell response can ultimately only be assessed by testing its functional properties. This has been extremely challenging in chronic HCV infection, but the availability of leukapheresis samples allowed the input of sufficiently large numbers of PBMCs to yield robust functional data (**Fig. 4a**). We tested for the secretion of IFN*γ*, TNF*α*, and IL-2 after *ex-vivo* peptide stimulation, as well as the mobilization of CD107A, a surrogate marker for T cell cytolytic capability. We found very limited cytokine secretion in HCV-specific T_EX_ pre-treatment, as described previously^35^, while a significant proportion of the cells displayed a cytotoxic response (**Fig. 4b-d**). These results were almost unchanged after HCV cure, apart from a significant, but low-level increase in TNF*α* secretion. In contrast, T_ESC_ cells were more polyfunctional and already produced significantly more of each cytokine before treatment, at overall levels similar to those of T_MEM_ from spontaneously recovered HCV infection (**Fig. 4b-d**). Overall, the data indicate that early loss of antigenic stimulation, after the development of viral escape mutations, can lead to functionally active cells, whereas viral cure after extended chronic viremia does not. Further, even when cells are similar in phenotype, they can be significantly different in function.

**Figure 4:**
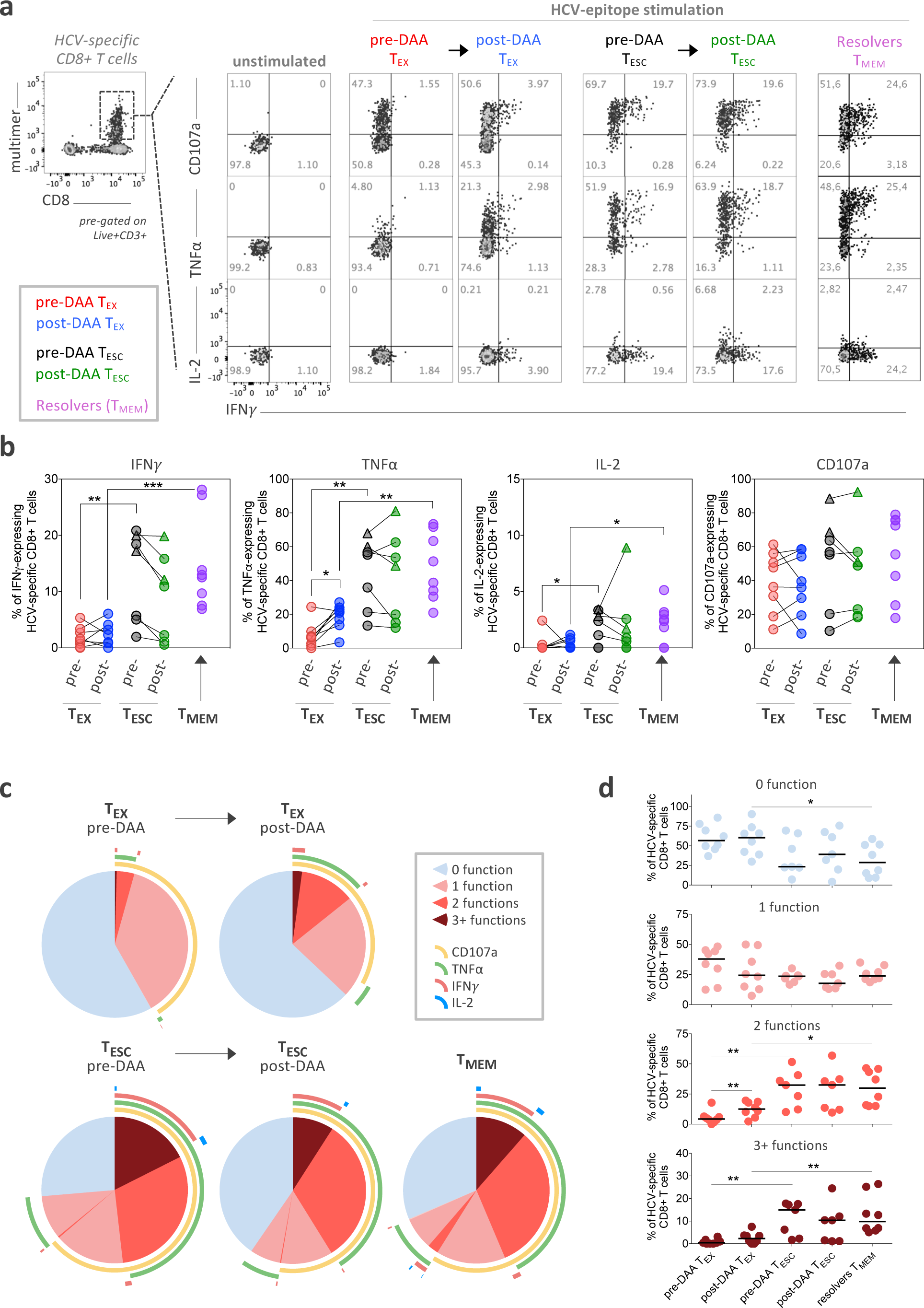
Functional analysis reveals that T_ESC_, but not T_EX_ after viral cure, display functional properties similar to those of T_MEM_ from HCV natural resolvers. **a**, Representative flow cytometry plots of the cytokine production and cytotoxicity capability of HCV-specific CD8+ T cells following *ex vivo* stimulation with or without cognate antigen. The co-expression patterns and percentage of cells producing IFN*γ*, TNF*α* and IL-2 cytokines and expressing CD107a are indicated. **b**, Dot plot histograms displaying IFN*γ*, TNF*α* and IL-2 production as well as CD107a expression across T_EX_ (paired samples, n=8) and T_ESC_ (paired samples, n=7) pre- and post-DAA therapy, and in resolver T_MEM_ (n=8). Triangles identify two analyzed T_P-ESC_ populations among the T_F-ESC_. Comparison rules and statistical tests have been performed as presented in Extended Data Fig.3c. **c**, Overlapping pie-charts describing the polyfunctionality of T_EX_ (paired samples, n=8) and T_ESC_ (paired samples, n=7), pre- and post-DAA therapy, as well as T_MEM_ (n=8), after *ex vivo* stimulation with cognate antigens. **d**, Frequencies of T cells with one, two and three or more functions as defined by the expression or co-expression of CD107a, IFN*γ*, TNF*α* and IL-2 after stimulation cognate antigens.

### The T_EX_ phenotypic and functional profile shows limited evolution with additional time post-HCV cure

To establish whether our assessment of the effect of HCV cure on T_EX_ at week 24 post treatment initiation might be missing important subsequent changes over time, we followed 4 patients with additional leukapheresis samples up to 3 years post initiation of treatment. We found that many molecules had already reached steady state expression levels at the early post-treatment timepoint, including CD38, HLA-DR and PD-1 (**Fig. 5a** and **Extended Data Fig. 4a**). Nevertheless, some markers continued to change beyond month 6, with a further increase of CD127 expression and additional decreases for Eomes and CD39 expression (**Fig. 5b** and **Extended Data Fig. 4a**). However, these changes had no significant impact on T cell activation and function, e.g. on CD69 and CD107a upregulation or IFN*γ* and TNF*α* production following cognate antigen stimulation, which all remained stably diminished compared to T_MEM_ (**Fig. 5c,d** and **Extended Data Fig. 4b**). Overall, the data do not indicate substantial improvement of T_EX_ during long-term observation after HCV cure, but suggest fixation of T_EX_ in their dysfunctional state after cessation of chronic antigen stimulation.

**Figure 5:**
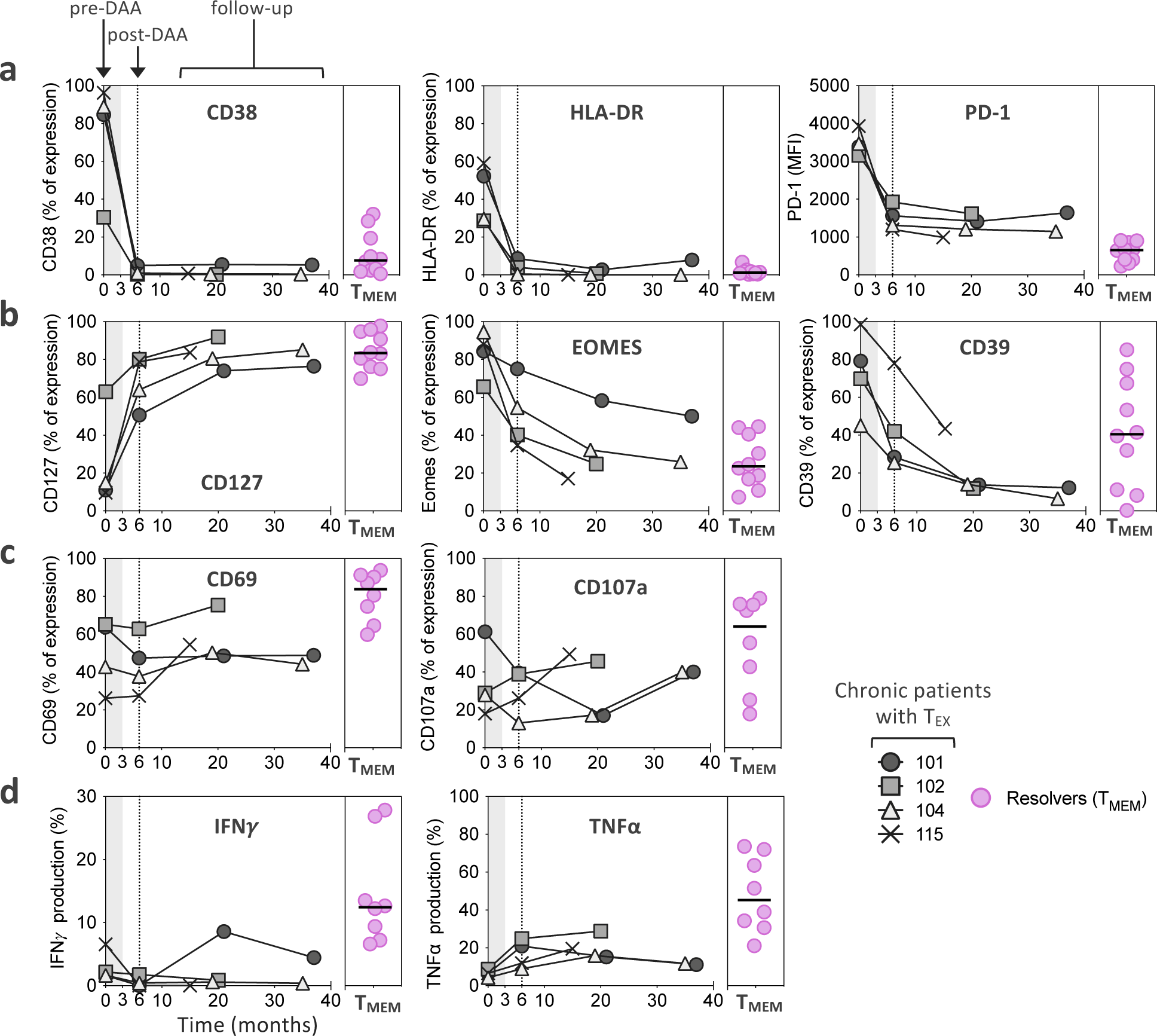
The T_EX_ phenotypic and functional profile shows limited evolution over time post-HCV cure. **a-b**, Temporal dynamics of expression levels of CD38, HLA-DR and PD-1 (**a**), as well as CD127, EOMES, and CD39 (**b**), by T_EX_ from four chronic HCV patients treated with DAA. **c-d**, Functional analysis of CD69 and CD107a (**c**), as well as IFN*γ* and TNF*α* production (**d**) after *ex vivo* stimulation with cognate antigen. Expression levels are displayed as percentage of expression except for PD-1, which is expressed as MFI. Expression levels in T_MEM_ from spontaneous resolvers are displayed for comparison.

### Transcriptional analysis of HCV-specific CD8+ T cells confirms broad changes in T_EX_ after removal of antigen, but also identifies scars in the transcriptional landscape

To broaden the analysis to all molecules expressed in antigen-specific CD8+ T cells, we performed RNAseq on sorted HCV multimer-binding CD8+ T cell populations for the previously analyzed T cell categories (T_EX_ and T_ESC_ pre- and post-DAA therapy, and T_MEM_). First, we compared gene expression in T_EX_ before and after HCV cure. Confirming our findings from flow-based phenotyping, we found broad changes in gene expression after antigen removal, with a total of 578 differentially expressed genes (**Fig. 6a** and **Extended Data Fig. 5a**). This included already detected changes towards a memory phenotype, such as upregulation of CD127, but many additional molecules were expressed at similar levels to T_MEM_ (**Extended Data Fig. 5b**). This phenotypic change is further corroborated by the observation that differential gene expression between T_MEM_ from resolvers and T_EX_ dropped from 585 genes pre DAA treatment to 217 genes post DAA treatment (**Fig. 6a**). Principal component analysis of these three populations accordingly showed a change from complete separation of T_EX_ to partial overlap with T_MEM_ post-treatment (**Fig. 6c**). The transcriptional data also confirmed on a much broader scale the more limited changes in T_ESC_ after therapy, with most changes in T_ESC_ also observed in T_EX,_ but with smaller differences between pre and post treatment (**Fig. 6b** and **Extended Data Fig. 5c**). The transcriptional data also show that T_ESC_ do not exhibit changes in key pathways such as PD-1 response signaling or in their memory phenotype after HCV cure, in contrast to T_EX_ (**Fig. 6d**). While T_ESC_ already had more memory-like characteristics before treatment, T_EX_ still had exhausted characteristics compared to T_MEM_ after antigen removal, including lower translation initiation and higher expression of the KLRG1-related gene cluster^5, 36^ (**Fig. 6e**). Together, these data confirm and expand the phenotyping analysis by flow cytometry, showing broad and significant changes towards memory in T_EX_ post cure, while in T_ESC,_ these changes have already occurred before, after viral escape.

**Figure 6:**
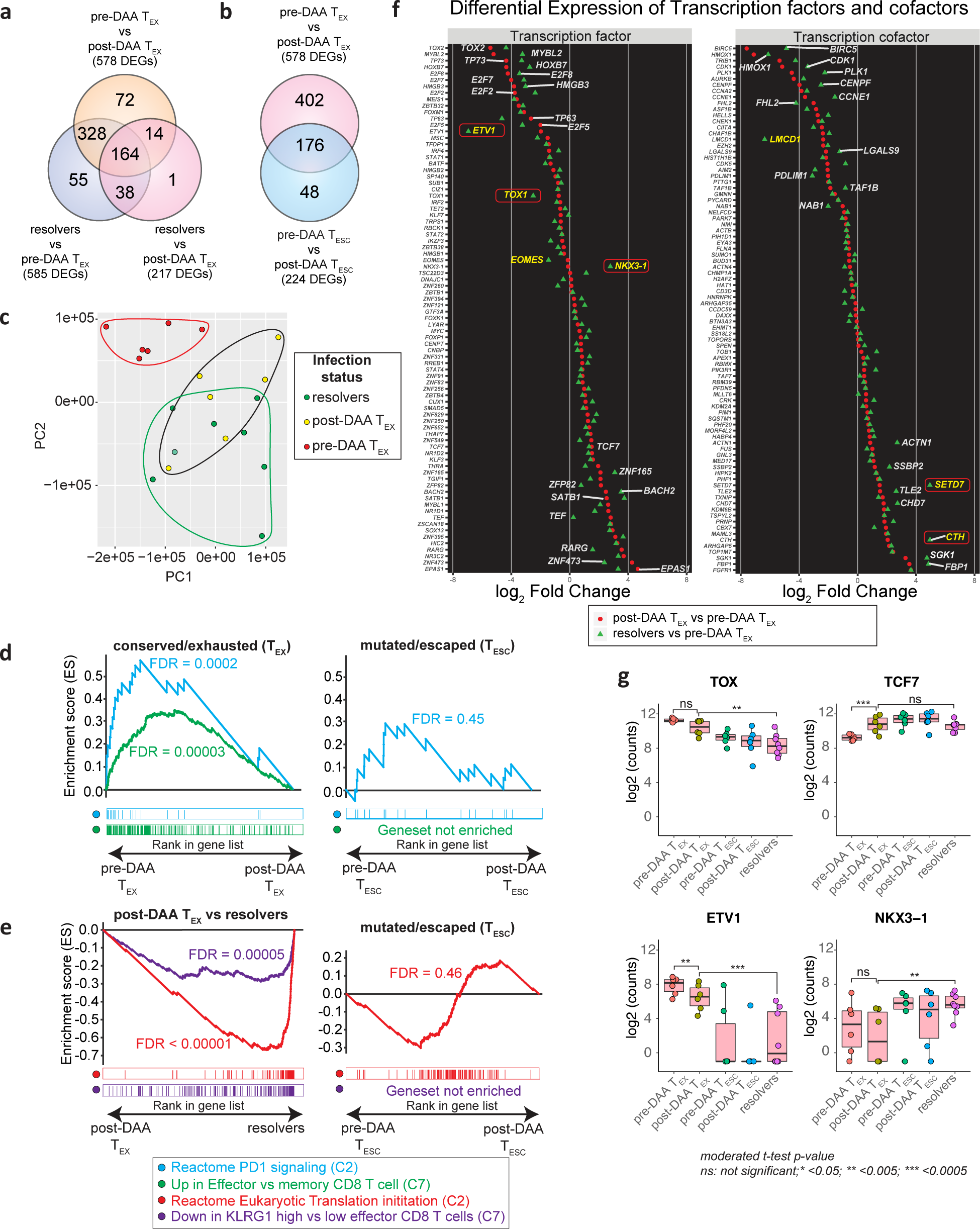
Transcriptional analysis confirms broad changes in T_EX_ after removal of antigen, but also identifies exhaustion scars in the transcriptional landscape. HCV-specific MHC-multimer-sorted CD8+ T cells from patients pre- and post-DAA treatment were compared with those from patients who resolved infection spontaneously. **a**, **b**, Venn diagrams showing the number of differentially expressed genes (DEGs) and overlap between T_EX_ pre- vs post-DAA therapy (paired samples, n=6), vs T_MEM_ from resolver patients (n=8) (**a**), or vs T_ESC_ (paired samples n=6) (**b**). **c**, Principal component analysis of gene expression profiles from pre- DAA T_EX_, post-DAA T_EX_, and resolver T_MEM_ cells. **d**, Gene set enrichment analysis (GSEA) of the transcriptional signature “reactome PD1 signaling (C2)” and “Up in effector vs memory CD8+ T cell (C7)” enriched in pre-DAA T_EX_ cells compared post-DAA (left panel) or in pre- DAA T_ESC_ cells compared post-DAA (right panel). **e**, GSEA of transcriptional signature “reactome translation (C2)” and “Down in KLRG1 high vs low effector CD8+ T cells (C7)” enriched negatively in resolver T_MEM_ cells compared to post-DAA T_EX_ cells (left panel) or in pre-DAA T_ESC_ cells compared to post-DAA (right panel). **f**, Plot showing the log_2_ fold change difference of transcription factors and cofactors between post-DAA T_EX_ and resolver T cells with respect to pre-DAA T_EX_ cells. Transcription factors and cofactors that did not recover are highlighted in yellow font and with a red frame if statistical validation was made on 9 additional T_EX_ populations post-DAA therapy and as compared to T_MEM_ (Extended Data Fig. 6c). **g**, Expression (log_2_ counts) of TOX, TCF7, ETV1 and NKX3-1 in T_EX_ and T_ESC_ pre- and post-DAA (paired samples n=6) as well as in resolver T_MEM_ cells (n=8).

Since phenotypic improvements were incomplete and did not lead to functional improvement, we searched for genes that remained fixed in a T_EX_ state after antigen removal (top 20 differentially expressed genes between T_EX_ post-treatment and T_MEM_ in **Extended Data Fig. 5d**). We focused on transcription factors and transcription co-factors, as we hypothesized these to be at the center of potential immunological scars that prevent differentiation into effective memory T cells. While the expression levels of many transcription factors and co-factors in T_EX_ changed after treatment to levels comparable with those in T_MEM_, some notable outliers either did not change at all or changed insufficiently (**Fig. 6f**). An important example is the transcription factor TOX, which was recently identified as a key driver of T cell exhaustion (**Fig. 6f,g**)^29, 37, 38^. TOX expression decreases post antigen removal but does not reach the level of either T_ESC_ or T_MEM_, in contrast to TCF7, which fully normalizes. The same pattern was observed for the transcription factor Eomes and additional transcription factors and co-factors, most prominently ETV-1, NKX3-1, LCMD-1, SETD7, and CTH (**Fig. 6f,g** and **Extended Data Fig. 6a**). Of these transcription factors and co-factors, ETV1, EOMES and LCMD1 displayed expression patterns correlating with TOX expression, suggesting potential mechanisms of coregulation (**Extended Data Fig. 6b**). We then validated the differential expression of these molecules between T_EX_ post-DAA treatment and T_MEM_ in 9 additional T_EX_ populations post therapy, confirming the gene expression differences for TOX, ETV1, NKX3-1, SETD7, and CTH, but not LMCD1. Eomes showed similar difference as in the original comparison, but this did not reach statistical significance (**Extended Data Fig. 6c**). Importantly, most of these molecules without clear recovery displayed much more T_MEM_-like expression levels in T_ESC_ (**Fig. 6g** and **Extended Data Fig. 6a**). These data suggest that, despite the broad transcriptional changes in T_EX_ post-DAA treatment, many key transcriptional regulators remain in a T_EX_ state and thus seem to represent immunological scars, which are mostly absent in T cells that had received antigen signal for a more limited time due to viral escape mutations.

## Discussion

T cell exhaustion is a key feature of chronic viral infections and cancer. As demonstrated by the success of checkpoint inhibitor therapies for a variety of malignant tumors, T cell exhaustion can be reversible. However, the efficacy of current checkpoint inhibitors is not universal, and recovery is usually not lasting. In addition, other approaches to resurrect T cell responses in chronic viral infections, such as therapeutic vaccines and immunomodulatory agents, will also depend on their ability to drive exhausted T cells out of their dysfunctional state. Thus, a better understanding of the molecular pathways underlying T cell exhaustion and its reversal is needed to develop improved therapeutic approaches. Here we utilized treatment of chronic HCV infection with DAA, the only current model of a cured chronic viral infection, to test the hypothesis that drug-induced complete clearance of chronic antigen stimulation allows T_EX_ to differentiate towards a memory profile without additional immunotherapeutic intervention. Early reports of DAA-cured chronic HCV infection indicated some reversal of T cell exhaustion, with increased proliferative capacity of HCV-specific CD8+ T cells^28^, lower expression of PD-1 and a shift towards a TCF-1+ CD127+ memory-like T cell phenotype^27^. However, these cells did not fully resemble T_MEM_^27, 29, 30^ and, at least in one patient with re-exposure to HCV due to viral relapse, these cells expanded while reverting to an exhausted T cell profile and were unable to contain viral replication^27^.

To define the impact of antigen removal on T_EX_ more broadly and in greater depth, we analyzed 20 subjects undergoing identical DAA treatment for chronic HCV infection in a clinical trial optimized for immunological studies. This study protocol included leukapheresis collection for at least 2 timepoints, one pre- and one post-treatment, allowing an integrated approach with flow cytometry-based phenotyping, functional studies, and RNAseq and TCR repertoire analyses, all on the same research samples. Phenotypically, we observed broad changes in T_EX_ post antigen removal, with expression levels of 23/37 parameters tested by flow cytometry changing significantly, all towards a memory phenotype. Importantly, this was not caused by selective emergence or disappearance of phenotypically distinct T cell clones, as we observed rather stable clonal repertoires between pre- and post-DAA T_EX_ despite the observation of many molecules changing expression levels in almost the complete T_EX_ population. Flow data were further expanded by RNAseq analysis, revealing a total of 578 differentially expressed genes between T_EX_ pre- and post-treatment. This finding demonstrates that the previously described changes post antigen removal, most notably the shift towards TCF-1+ CD127+ memory-like T cells^27^, while critical, are just one component of a broadly altered transcriptional landscape in T_EX_ after cure. Importantly, our data also suggest that the observed changes are mostly a function of terminated TCR signaling, and not mediated by the resolution of the chronic inflammatory environment, since T cells against other viruses remained unchanged in their phenotype and those targeting HCV escape variants displayed much more limited changes on both the protein and transcriptional level.

Through comparison of the T_EX_ post antigen removal with bona fide T_MEM_ from patients who had spontaneously resolved infection, it became apparent that the differentiation of T_EX_ towards memory was incomplete. Significant differences in gene expression remained for about one third of the molecules that were differentially expressed between T_MEM_ and T_EX_ before treatment, including key mediators of T cell exhaustion such as PD-1 or TOX. While the number of genes that did not recover was about half the number of those that did, functional analysis revealed that the fixed or scarred genes must be especially relevant, as the broad transcriptional changes in T_EX_ post treatment failed to translate into a meaningful increase in cytokine production. Additionally, we found that T cells targeting escaped HCV epitopes could be functionally as robust as T_MEM_ from spontaneously resolved infection, even before treatment, even though their phenotype was relatively close to post-treatment T_EX_. This observation highlights that even the broadest and most detailed phenotyping is insufficient to deduce T cell functionality. In the context of this observation, it is also important to consider that viral escape from HCV-specific CD8+ T cell responses almost exclusively occurs early in chronic infection, typically between 3 and 9 months after viral exposure^32, 33^. Thus, our results suggest that, after a short period of antigen exposure during early infection, T_ESC_ can still recover by antigen removal alone, in contrast to T_EX_ with many years of chronic antigen stimulation. An alternative explanation is that T cells targeting exhausted epitopes had a much longer time to recover, i.e. several years compared to the 23 weeks post-treatment mediated viral resolution in T_EX_. We deem this unlikely, as we observed limited additional changes in phenotype and no sign of functional recovery when we followed a subgroup of chronic patients for up to 3 years after treatment. The idea that duration of antigen stimulation, rather than duration of recovery, is the defining factor for the ability to differentiate into T_MEM_ is also supported by data from the LCMV model of viral infection, where early transfer of exhausted T cells into uninfected mice was similarly associated with improved T cell function, in contrast to transfer at later timepoints^8^.

Our results raise several important questions for future investigations. It is important to determine whether the global changes we observed in the T_EX_ population after antigen removal are based on re-differentiation of all T_EX_, or whether they are a consequence of preferred outgrowth of the pre-existing TCF-1+ memory-like population^39^. Our TCR analysis found a mostly preserved TCR composition of T_EX_ after DAA cure, demonstrating that the observed changes are not the consequence of a remodeling of the overall clonal repertoire. Together with data from a recent scRNAseq study suggesting that the HCV-specific TCF-1+ memory-like population represents the likely precursors to all terminally differentiated T_EX_^30^, it seems that chronic antigen removal after post HCV cure leads to loss of the terminally differentiated T_EX_ population within each clone. What seems to remain are the memory-like precursors that are nevertheless functionally impaired. Final proof for this hypothesis will have to come from more definitive mouse transfer experiments of the different T_EX_ subsets. Furthermore, some of the transcription factors we identified with similar expression behavior to that of TOX should be investigated further. To our knowledge, several of these molecules were not previously associated with T cell exhaustion. With most comprehensive exhaustion data coming from mouse models, this discrepancy could potentially be due to these molecules having different roles in mice and humans. Finally, it is critical to understand the root cause for the lack of plasticity in T_EX_, with a major candidate being epigenetic scarring preventing actual recovery of exhausted T cells. It will also be important to further establish whether there is a window of opportunity to correct or prevent this state early in T cell exhaustion. To this end, we are currently studying whether early DAA therapy during the first year of HCV infection leads to more profound reprogramming of exhausted T cell responses.

In summary, our study revealed a broad impact of antigen removal on human T_EX_ that did not translate into functional recovery. We also identified genes that seem irrevocably fixed in their expression levels after long-term antigen exposure, whereas T cells with a shorter duration of TCR stimulation seem more open to complete differentiation towards functional memory. These results suggest a window of opportunity for reconstituting immunity early in exhaustion and identify target genes for interventions to rescue more terminally exhausted CD8+ T cells.

## Materials and methods

### Study design, DAA treatment and sample collection

Patients with chronic genotype (GT) 1a HCV infection were enrolled in an open-label Phase 3 clinical trial of paritaprevir/ritonavir, ombitasvir, dasabuvir and ribavirin designed to evaluate the effect of successful antiviral therapy on innate and adaptive immune responses (NCT02476617)^31^. The clinical trial was approved by the Partners Healthcare Institutional Review Board and has been previously described^31^.

Patients were all chronically infected with HCV GT 1a and presented no additional co-infection such as HIV or HBV, or had previously resolved HCV infection on their own. Trial participants were treated for 12 weeks with a combination of paritaprevir/ritonavir/ombitasvir + dasabuvir + ribavirin and regular blood draws were collected pre- and post-DAA therapy, including at least two leukapheresis collections at weeks 0 and 24. Additional leukaphereses were collected from four trial subjects between weeks 60 and 80. Peripheral blood mononuclear cells (PBMCs) were extracted by Ficoll-Paque (GE Healthcare Life Sciences) density gradient centrifugation and frozen down for further processing, together with plasma samples. Repository-frozen PBMCs from an additional cohort of patients who had spontaneously resolved HCV infection (Resolvers) were also studied. Clinical characteristics of the patients are described in **Extended Data Table 1**. All patients provided written informed consent.

### Screening for *ex vivo* detection of virus-specific CD8+ T cell populations and cell enrichment

PBMC samples were thawed and analyzed by flow cytometry to identify virus-specific CD8+ T cell responses using appropriate HLA class I multimers, pre- and post-DAA treatment (**Extended Data Fig. 1a,b**). Magnetic bead enrichment of multimer-positive cells using anti-PE or -APC beads and LS columns (Miltenyi) was performed on up to 200 million PBMCs, according to the manufacturer’s instructions, to capture HCV-specific CD8+ T cell populations of even extremely low-frequency (**Extended Data Fig. 1c**).

### Sequencing HCV epitopes and testing of variant epitopes

We deep sequenced each of the identified HCV epitopes and tested the T cell recognition of epitope variants. HCV epitopes from chronically infected patients were sequenced from circulating viruses in plasma samples collected before trial initiation to identify potential escape mutations. An amplicon surrounding epitopes C63B, 127D, 140G, A2-198 and 174D was generated using the following conditions. The reaction consisted of 2x First Strand Buffer, sense (A2F: AAC GTT GCG ATC TGG AAG AC or A3F: GCT CTC ATG ACC GGC TTT AC) and antisense primers (A2R: GGA AGC GTG GTT GTC TCA AT or A3R: AGA GAT CTC CCG CTC ATC CT) at 0.4µM, and a Superscript III RT/Platinum *Taq* Mix (Invitrogen), with the following conditions: cDNA synthesis for 30 minutes at 50°C, followed by heat denaturation at 94°C for 2 minutes, the PCR amplification conditions were 40 x (94°C, 15 s; 55°C, 30 s; 68°C 180 s), with a final extension at 68°C for 5 minutes. For epitope 4H, an amplicon was generated using 2x First Strand Buffer, sense (UTR4: CCT TGT GGT ACT GCC TGA TAG) and antisense primers (A1R-1a: GGG YAG CAG TTG ACA CRA TCT) at 0.4µM, and a Superscript III RT/Platinum *Taq* Mix (Invitrogen), with the following conditions: cDNA synthesis for 30 minutes at 55°C, followed by heat denaturation at 94°C for 2 minutes, the PCR amplification conditions were 40 x (94°C, 15 s; 58°C, 30 s; 68°C 240 s), with a final extension at 68°C for 10 minutes. Finally, for epitopes 262G and 2226D the primers used were (A4F: CTC ACT GAT CCC TCC CAC AT) and antisense primers (A4R: GGG GAG GAG GTA GAT GCC TA) with the following PCR conditions 40 x (94°C, 15 s; 64°C, 30 s; 68°C 180 s), with a final extension at 68°C for 10 minutes. PCR products were visualized on a 1% agarose gel and purified using the PureLink Quick Gel Extraction Kit (Invitrogen). PCR amplicons were fragmented and barcoded using NexteraXT DNA Library Prep Kit, as per manufacturer’s protocol. Samples were pooled and sequenced on an Illumina MiSeq platform, using a 2 x 250 bp V2 reagent kit. Paired-end reads obtained from Illumina MiSeq were assembled into a *de novo* HCV consensus sequence and aligned with intra-host variants called as previously reported^40, 41^. Data has been deposited to the NCBI Sequence Read Archive under accession numbers SRR11811484 – SRR11811504. Wild-type and variant epitope sequences are available in **Extended Data Fig. 2a**. If the circulating virus contained epitope variants, we generated T cell lines by stimulating the patient’s PBMCs for 14 days with the wild-type peptide under addition of IL-2 and subsequently tested recognition of both wild-type and variant epitope by intracellular cytokine staining aiming to detect IFN*γ* production. The effect of epitope sequence variations on T cell recognition based on IFN*γ* production compared to wild-type epitopes was tested as previously described^23^, and T cell responses were categorized into two categories (**Extended Data Fig. 2b**): 1) partial recognition of the epitope variant (partially escaped T cells, T_P-ESC_) and 2) complete loss of T cell stimulation (fully escaped T cells, T_F-ESC_).

### *Ex vivo* immunophenotyping of antigen-specific CD8+ T cells by flow cytometry

To profile the different immune subsets shown in **Fig. 1c; Fig. 2a-f and Fig. 3**, surface staining and intracellular stainings of *ex vivo* multimer-positive T cells were performed. In brief, PBMC were thawed and washed in R10 medium [RPMI 1640 (Sigma-Aldrich), 2% fetal calf serum (FCS), 1.5% 1 M HEPES buffer (Fisher Scientific), 1× l-glutamine (Fisher Scientific) and 1× streptomycin–penicillin (Fisher Scientific)]. Cells were then stained with LIVE/DEAD Cell Viability dye (Thermo Fisher Scientific) according to the manufacturer’s protocol. The cells were washed in FACS buffer and incubated with select MHC class I Pentamers (ProImmune) or Tetramers (NIH tetramer core facility) for 10 minutes at room temperature. After a wash, cells were stained with surface antibodies for 30 minutes, washed twice, and a fixation– permeabilization step was performed using the FOXP3 Transcription Factor Staining buffer set (Thermo Fisher Scientific) according to the manufacturer’s protocol. Alternatively, multimer-positive cell enrichment was performed before the surface antibody staining step. Cells were then stained with intracellular antibodies. A full list of the antibodies used is available in **Extended Data Table 2**. All steps were performed at 4 °C unless otherwise specified by the manufacturer. Acquisition was performed on an LSR-II flow cytometer (Becton Dickinson).

### T cell stimulation with cognate antigen and ICS assays

For *ex vivo* analyses (for **Fig. 4** and **Fig. 5**), frozen PBMC samples were rested overnight at 37 °C in R10 medium [RPMI 1640 (Sigma-Aldrich), 2% FCS, 1.5% 1 M HEPES buffer (Fisher Scientific), 100× l-glutamine (Fisher Scientific) and 50× streptomycin–penicillin (Fisher Scientific)]. Rested PBMCs containing *ex vivo* antigen-specific CD8+ T cell populations, or alternatively previously generated T cell lines (for **Extended Data Fig 2b**), were then incubated with select MHC class I Pentamers (ProImmune) or Tetramers (NIH Tetramer Core Facility) for 10 minutes at room temperature before stimulation with or without cognate antigen at 10 μg/mL in the presence of 10 μL/mL anti-CD28/49d (BD Biosciences) and 1x protein transport inhibitor (eBioscience) at a final volume of 500 μL for 3 h. Additionally, 5 μL of anti-CD107a antibody was added to the stimulation mixture. Alternatively, multimer-positive cell enrichment was performed on rested PBMCs before cell stimulation. Following incubation, cells were washed and stained with a viability dye (LIVE/DEAD Fixable Blue; Thermo Fisher Scientific) according to the manufacturer’s protocol, and then stained with the surface antibodies followed by intracellular antibodies as described in the above section. A full list of the antibodies used is available in **Extended Data Table 2**. All steps were performed at 4 °C unless otherwise specified by the manufacturer. Acquisition was performed using an LSR-II flow cytometer (Becton Dickinson).

### Flow cytometry data analysis and visualization

Flow-cytometry data were analyzed using FlowJo software. Protein expression levels were extracted as percentages and/or as median fluorescence intensity (MFI) numerical values. Alternatively, flow-cytometry data were analyzed using Cytobank^42^ to generate the T-distributed stochastic neighbor embedding (t-SNE) visualization presented in **Fig.3e**, based on the expression levels of CD38, HLA-DR, PD-1, CD39, TIGIT, CCR7, CD45RA, Integrin-Beta-7 and CD62L. Extracted numerical values referring to the expression levels of the 37 proteins presented in **Fig. 2, 3** and **Extended Data Fig. 3** were integrated into Radar Charts (**Fig. 2 and 3**) using Excel software or into Principal Component analysis using R software (**Fig. 3f** and **Extended Data Fig. 3e**) or ClustVis^43^ (Extended Data **Fig. 3d**). Dot plot histograms integrating the expression levels of the 37 proteins across the different T cell populations analyzed were generated using GraphPad Prism software (**Extended Data Fig. 3b**).

### Cell sorting and RNAseq library preparation

*Ex vivo* HCV-specific CD8+ T cells were sorted from chronically infected patients pre- and post-DAA therapy (T_EX_, paired samples, n=6; T_ESC_, paired samples, n=6), from patients with spontaneously resolved HCV infection (T_MEM_, n=6), as well as from a validation cohort of additional chronic patients post-DAA therapy (T_EX_, n=9) using the gating strategy described in **Extended Data Fig. 3a** on a FACS ARIA II sorter (BD Biosciences). Cells were sorted on RLT+, 1% BME buffer and were immediately frozen. RNA was extracted using RNeasy Micro Kit (QIAGEN) according to manufacturer’s instructions, treated with DNase I (New England Biolabs), then concentrated using Agencourt RNAClean XP beads (Beckman Coulter). Full-length cDNA and sequencing libraries were prepared using the Smart-Seq2 protocol as previously described^44^. Libraries were loaded on a Novaseq 6000 (Illumina) or on a Nextseq 500 (Illumina) to generate 150 base pair or 38 base pair, paired-end reads, respectively. Raw sequencing reads were aligned to the hg38 (GENCODE [v32]) reference transcriptome using STAR (v2.7.3a). Gene expression was quantified using RSEM (v1.3.1).

### TCR sequencing and analysis

HCV specific CD8 T cells from chronically infected patients pre- and post-treatment were sorted by flow cytometry into 500uL of Hepes Buffer (PBS+2%FBS+0.025M Hepes). After sorting, cells were pelleted and snap frozen, followed by DNA extraction using the Qiagen QIAmp DNA micro kit. Samples were eluted in 100uL of AE buffer and DNA concentration measured by nanodrop. Sample concentration was adjusted to 11 to 14ng/µl in a total volume of 36ml as recommended by the manufacturer. DNA amplification and library preparation were performed using the immunoseq hsTCRB kit from Adaptive Biotechnologies, with each DNA sample being amplified in duplicate. We also ran two negative control samples containing only AE buffer. The immunoseq assay is a multiplex PCR-based method that amplifies rearranged TCR CDR3 sequences and characterizes tens of thousands of TCRB CDR3 chains simultaneously. The assay captures the full TCR repertoire including specific individual clones and provides a method to identify and track common and rare clones. After library preparation with the immunoseq kit, quality control of the library was performed using Tapestation and qPCR before sequencing. The library was sequenced by the Biopolymers Facility of Harvard Medical School using Illuminaseq Next Gen500. Raw sequencing data were processed by Adaptive Biotechnologies using a multistep procedure that includes annotation, nucleated cell and TCRB quantification, and quality assessment to generate a comprehensive and quantitative report of each sample’s T-cell receptor repertoire.

### Differential gene expression analysis

Genes with non-zero counts were taken for *limma* analysis (linear models for microarray data). RNAseq count data were normalized using trimmed means of M-values (TMM) with the *calcNormFactors* function in *edgeR* package^45^. *Voom* was used to estimate the mean-variance relationship of log_2_ counts and precision weights were generated before linear modeling^46^. To identify differentially expressed genes (DEGs) between HCV-specific T cell populations in different disease outcomes, we performed multi-group analysis in *limma*^47^. Comparisons used exhausted T cells (pre-DAA and post-DAA), escaped T cells (pre-DAA and post-DAA), central memory T cells (pre-DAA, post-DAA and resolvers) and specific T cells from resolvers (complete results in **Extended Data Table 3**). DEGs between two groups were calculated using the moderated t-test based on an empirical Bayesian algorithm to estimate changes in gene expression (implemented using R package version 3.6.3; *limma* package version 3.42.2)^47^. False discoveries occurring due to simultaneous testing of the hypothesis were adjusted by applying the Benjamini-Hochberg procedure (adjusted P value)^48^. An adjusted P-value of 0.05 and absolute log_2_ fold change of 1 were considered cut-offs to create the DEG list using the *TopTable* function in the *limma* package. Principal component analysis was done using the *prcomp* function in R using normalized counts. Heatmaps for normalized counts were plotted using the *heatmap.2* function in the *gplot* R package. Gene counts of exhausted T cells (post-DAA) from the validation cohort (n=9) were compared with resolver samples from the original cohort (n=8) after batch correction using COMBAT^49^. For calculating correlations between genes, we used Pearson correlation.

### Gene set enrichment analysis (GSEA)

DEGs for each comparison (multi-group adjusted p-value of 0.05 as cut-off) were ranked based on their log_2_ fold change. GSEA was performed on ranked gene lists using the PreRanked method in the GSEA desktop application (version 4.0.0)^50, 51^. Enrichment scores were normalized by multiple sample correction to generate a normalized enrichment score (NES).

### Transcription factor analysis

We extracted genes that were significantly differentially expressed (adjusted p-value cut-off of 0.05) and identified transcription factor and cofactor genes among them by matching them to a list generated based on various databases^52, 53^.

### Statistical analyses

Statistical analyses were performed using GraphPad PRISM or R software. Statistical tests were used as indicated. *P < 0.05, **P < 0.01, ***P < 0.001, ****P < 0.0001.

## ACKNOWLEDGMENTS

We are grateful to the patients participating in the clinical trial. Important contributions were made by the HSCI-CRM Flow Cytometry Core Facility at MGH, the NIH Tetramer Core Facility, and the University of Oklahoma Medical Center HLA Typing Facility (William Hildebrand). We also thank all members of the Lauer Lab for insightful comments, critical reading of the manuscript and advice on figure design. This work was supported by NIH grants U19 AI086230, U01 AI131314 and R01 DA046277, and AbbVie sponsored the clinical trial (NCT02476617).

## AUTHOR CONTRIBUTIONS

P.T., D.W., and G.M.L. conceived and designed the experiments.

P.T., D.W., J.A.J., R.C.H., M.D., H.D., L.B., D.C.T., D.J. B., A.T.C., M.R., D.K., N.A., A.C., J.A.C., L.M., and T.E. performed and analyzed experiments.

D.W., S.S., D.L., and D.R.S. analyzed RNAseq data.

R.T.C., A.Y.K., and G.M.L. designed the clinical trial and patient selection.

J.B., J.G., L.L.L., and J.A. contributed to the clinical cohort recruitment and clinical database management.

G.M.L., N.H., and W.N.H. supervised RNAseq experiments and data analysis.

T.M.A. designed and supervised the viral sequencing study.

P.T., S.S., D.S., and G.M.L. drafted the manuscript with the help of all other authors.

---

**Extended Data Fig. 1:**
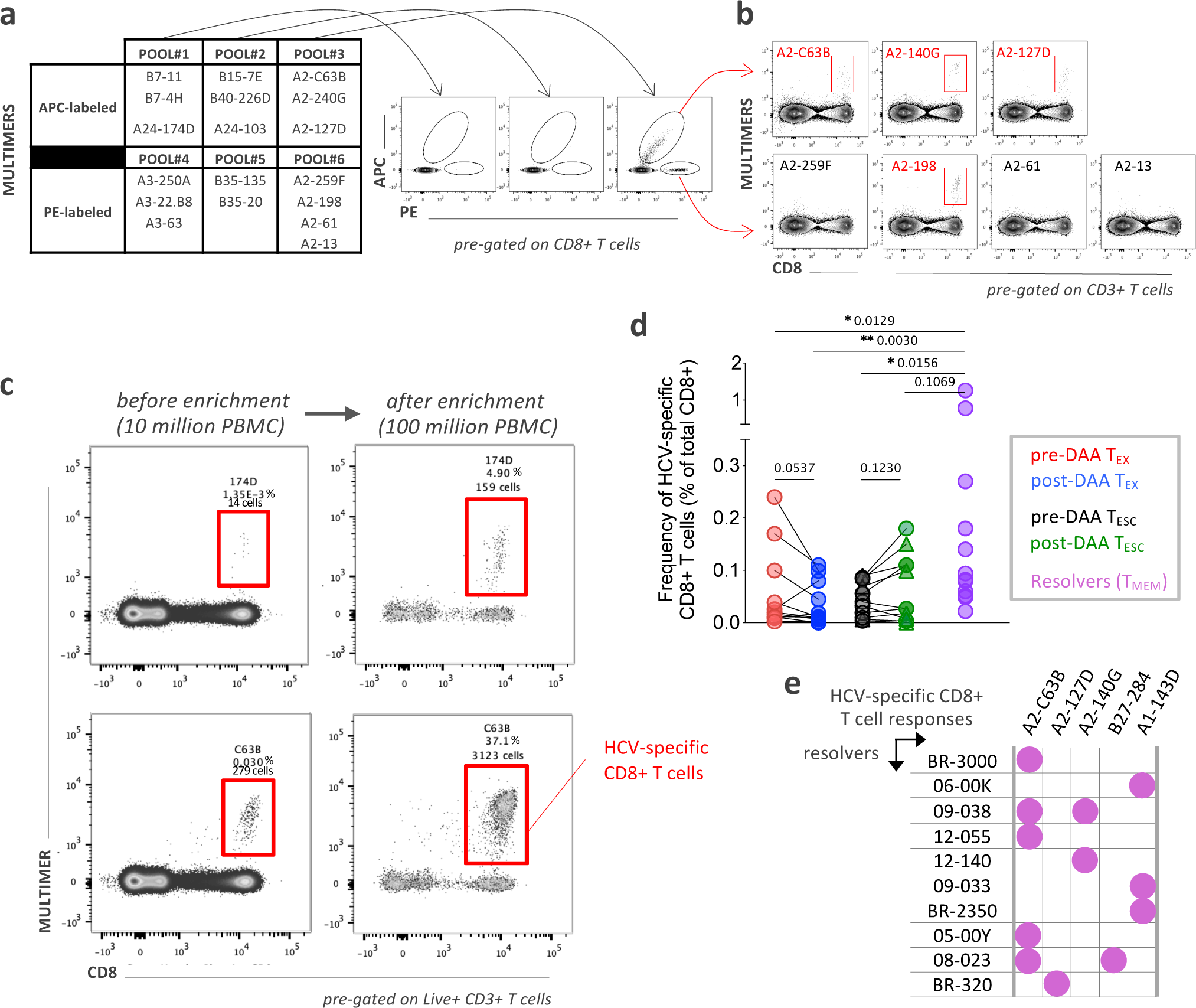
**a**, Screening strategy to detect HCV-specific CD8+ T cells by flow cytometry using pools of Class I MHC multimers labeled with PE or APC. **b**, Positive detection was followed by individual multimer staining to identify and/or distinguish multiple HCV-specific CD8+ T cell responses. **c**, Representative flow cytometry dot plots of HCV-specific CD8+ T cells before and after magnetic bead enrichment. **d**, HCV-specific-multimer positive CD8+ T cell frequencies, pre- and post-DAA treatment or after spontaneous resolution of the infection. Statistical testing by Wilcoxon tests (paired, nonparametric, for T_EX_ and T_ESC_ pre- and post-DAA) or Mann-Whitney tests (unpaired, nonparametric, when compared to Resolvers). **e**, HCV-specific CD8+ T cell responses (recognized epitopes and associated HLA class I restrictions) detected in a cohort of patients with spontaneously resolved HCV infection.

**Extended Data Fig. 2:**
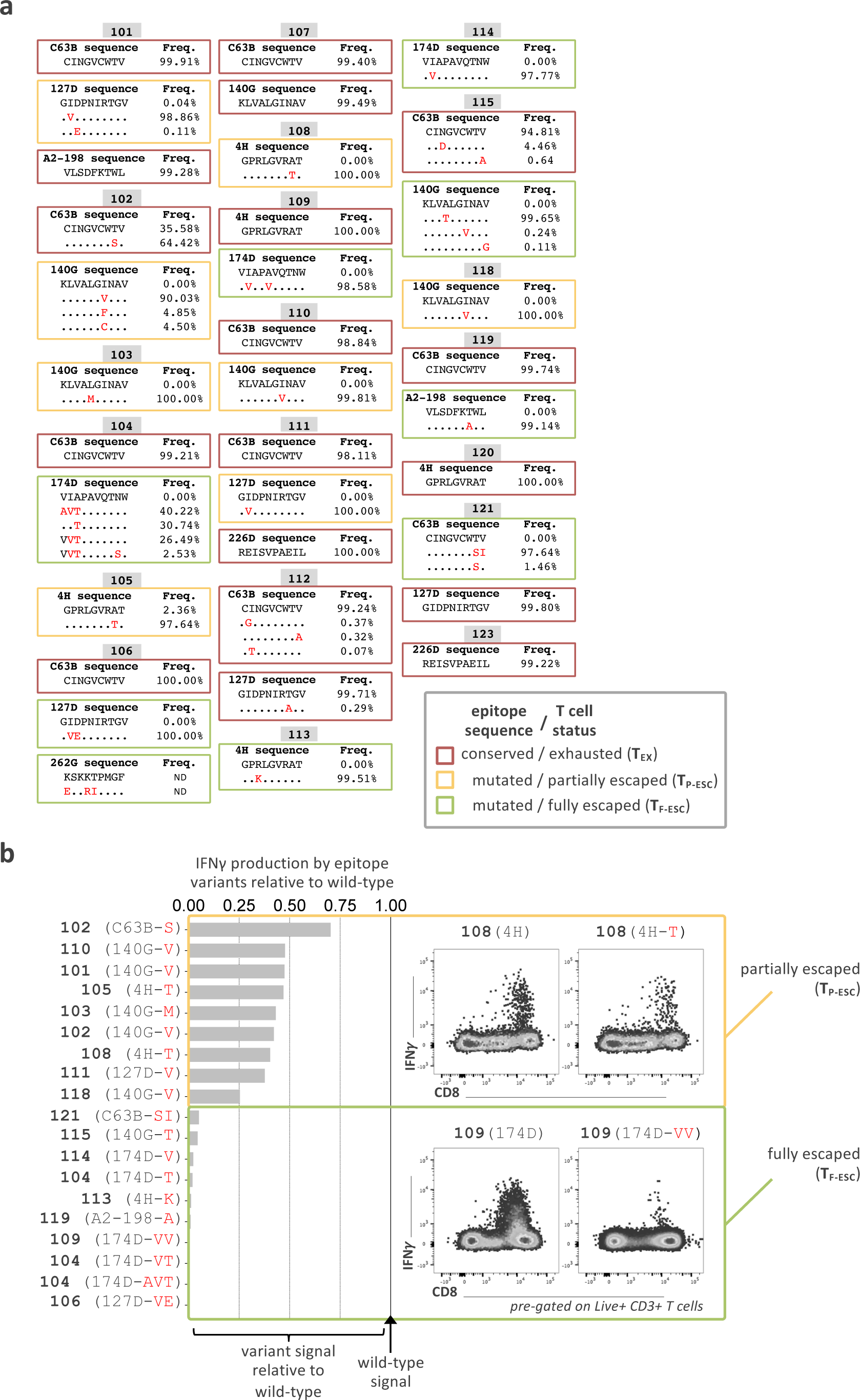
**a**, Sequences of select HCV epitopes across patients included in this study. Escape mutations are written in red and T cell status, whether the cells have full recognition of the virus (T_EX_) or are partially (T_P-ESC_) or fully escaped (T_F-ESC_) is indicated as determined through functional assays of the recognition of the variant epitopes compared to wild-types by intracellular cytokine detection of IFN*γ* by flow-cytometry (**b**).

**Extended Data Fig. 3:**
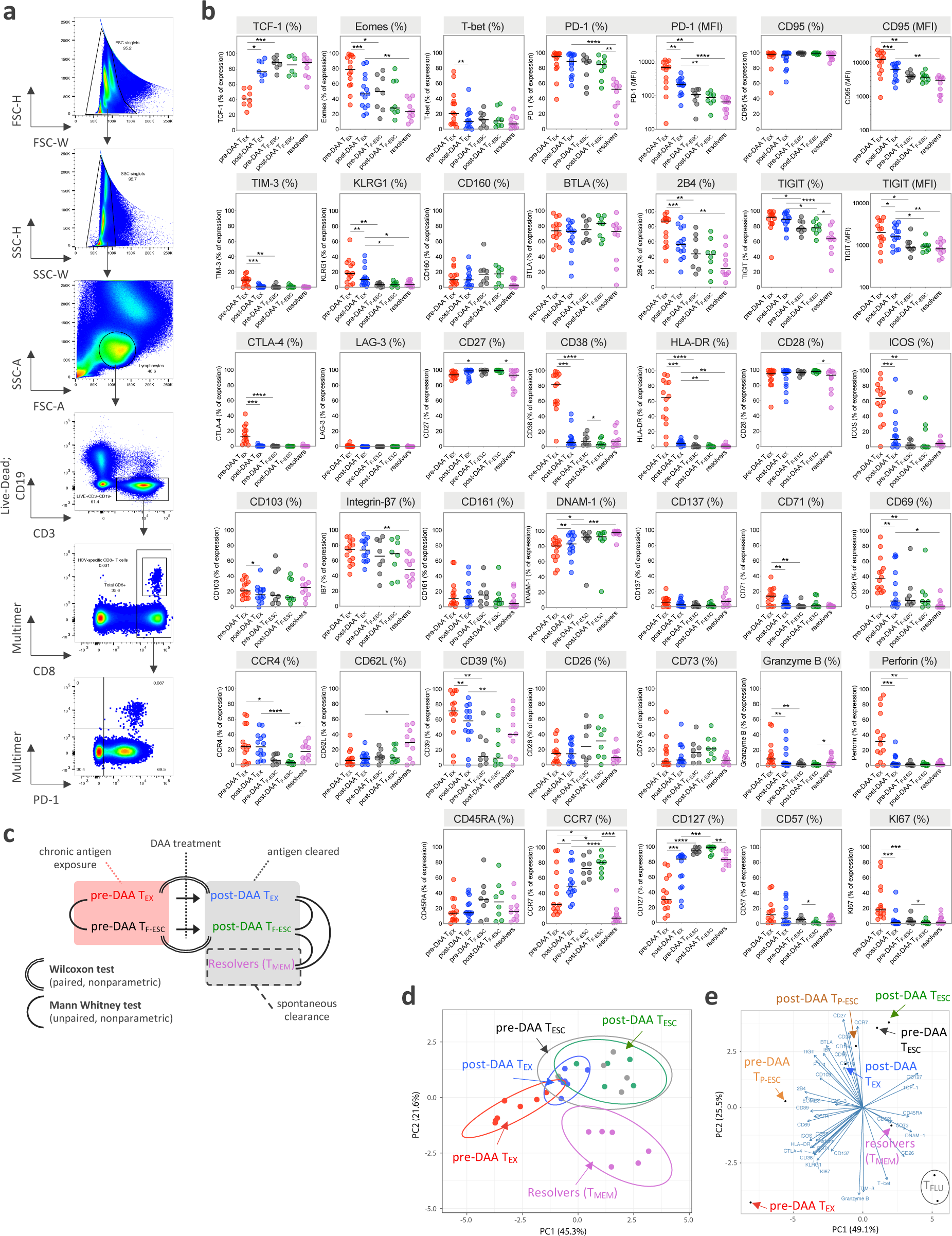
**a**, Flow cytometry gating strategy and representative flow cytometry dot plots. **b**, Dot plot histograms displaying the expression levels of the 37 proteins analyzed by flow cytometry across T_EX_ and T_F-ESC_, pre- and post-DAA therapy, and in resolver T_MEM_. **c**, Schematic representation of the comparison rules and statistical tests used to compare expression levels across the different T cell populations of interest. **d**, Principal component analysis of T_EX_ and T_F-ESC_, pre- and post-DAA therapy, as well as resolver T_MEM_, based on the expression levels of CD38, HLA-DR, PD-1, CD39, TIGIT, CCR7, CD45RA, Integrin-Beta-7 and CD62L, and as presented also in Fig. 3e by t-SNE analysis. **e**, Principal component analysis based on the expression levels of the 37 proteins analyzed by flow-cytometry and expressed by T_EX_, T_P-ESC_, T_F-ESC_ and T_FLU_, pre- and post-DAA therapy, as well as by resolver T_MEM_, with respective contribution and direction (arrows) of each of the 37 different proteins throughout PC1 and PC2 dimensions.

**Extended Data Fig. 4:**
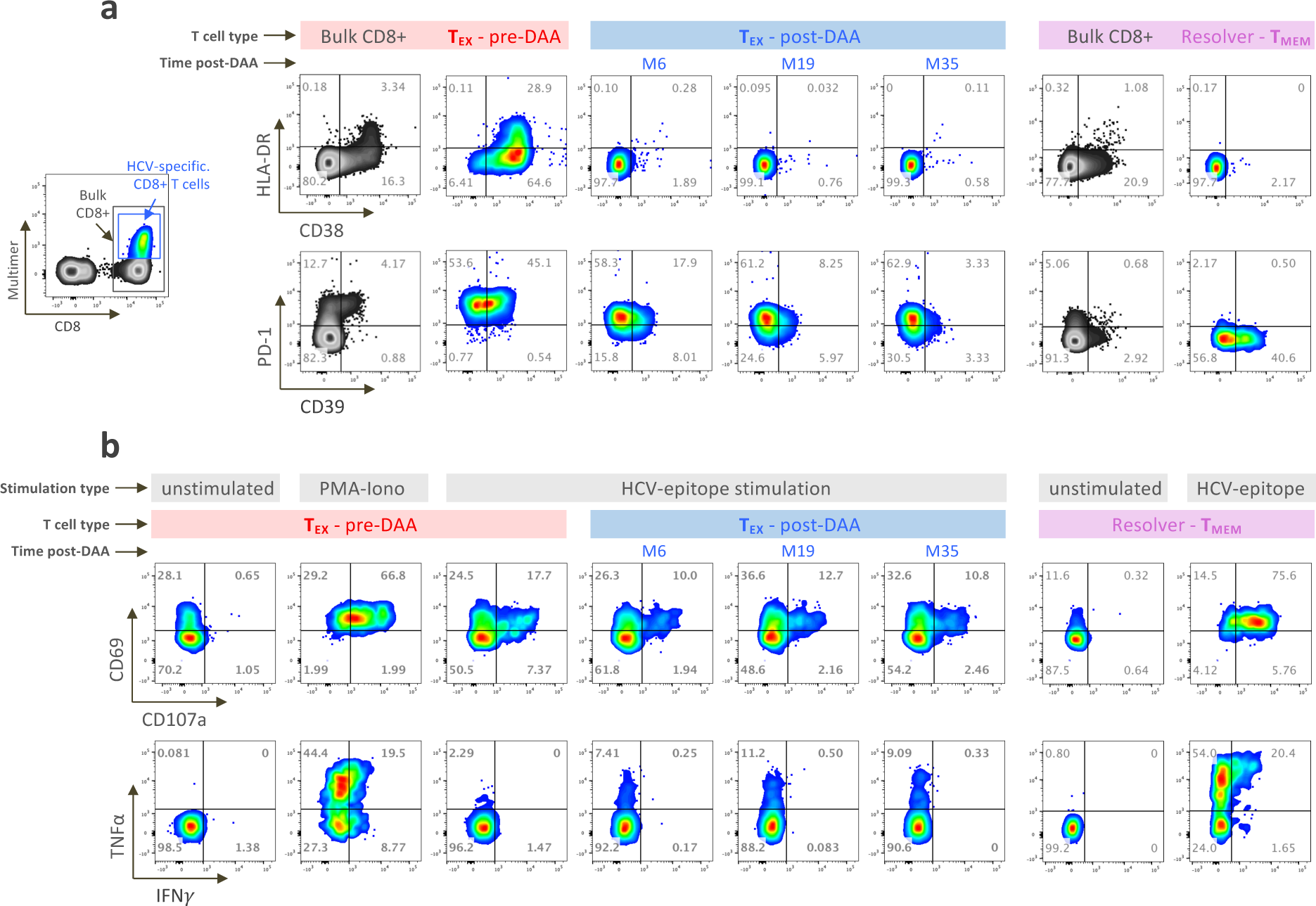
**a**, Representative flow cytometry dot plots showing the expression and co-expression patterns of HLA-DR and CD38 (upper panels) as well as PD-1 and CD39 (lower panels) by bulk CD8+ T cells (grey dots) or HCV-specific CD8+ T cells (colored dots), pre- and overtime post-DAA therapy or after spontaneous resolution. **b**, Representative flow cytometry plots showing the expression and co-expression patterns of CD69 and CD107a (upper panels) as well as IFN*γ* and TNF*α* cytokines (lower panels) by HCV-specific CD8+ T cells following *ex vivo* stimulation with or without cognate antigens, pre- and overtime post-DAA therapy or after spontaneous resolution.

**Extended Data Fig. 5:**
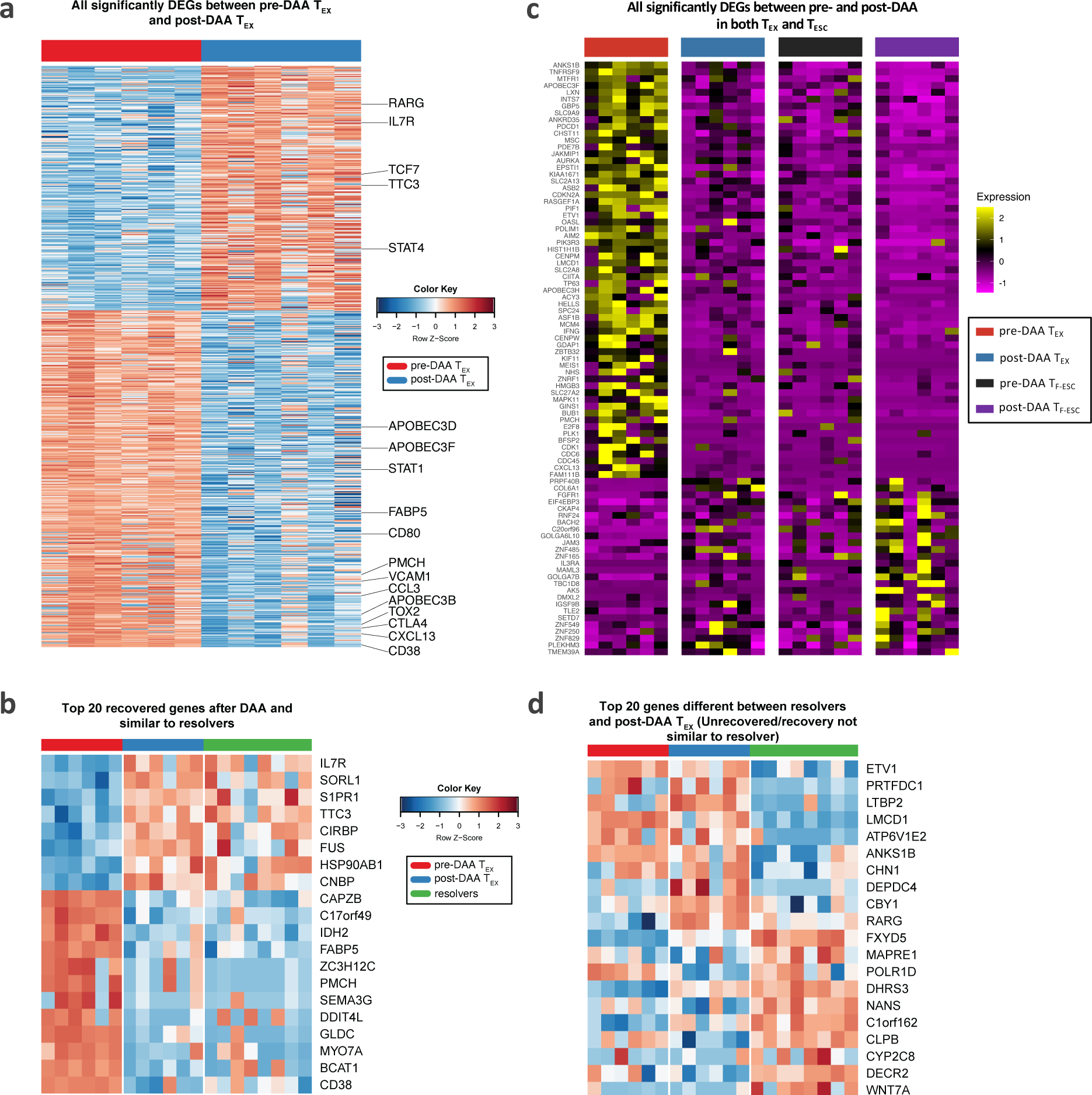
**a**, Heatmap showing all genes that were differentially expressed between pre-DAA T_EX_ and post-DAA T_EX_. **b**, Top 20 recovered genes after DAA treatment which are similar to resolver T cells. **c**, Heatmap showing the 176 significantly DEGs between pre- and post-DAA that are shared by T_EX_ and T_ESC_, as described in Fig. 6b. **d**, Heatmap showing the top 20 unrecovered genes after DAA treatment which were significantly different between post-DAA T_EX_ and T_MEM_ cells.

**Extended Data Fig. 6:**
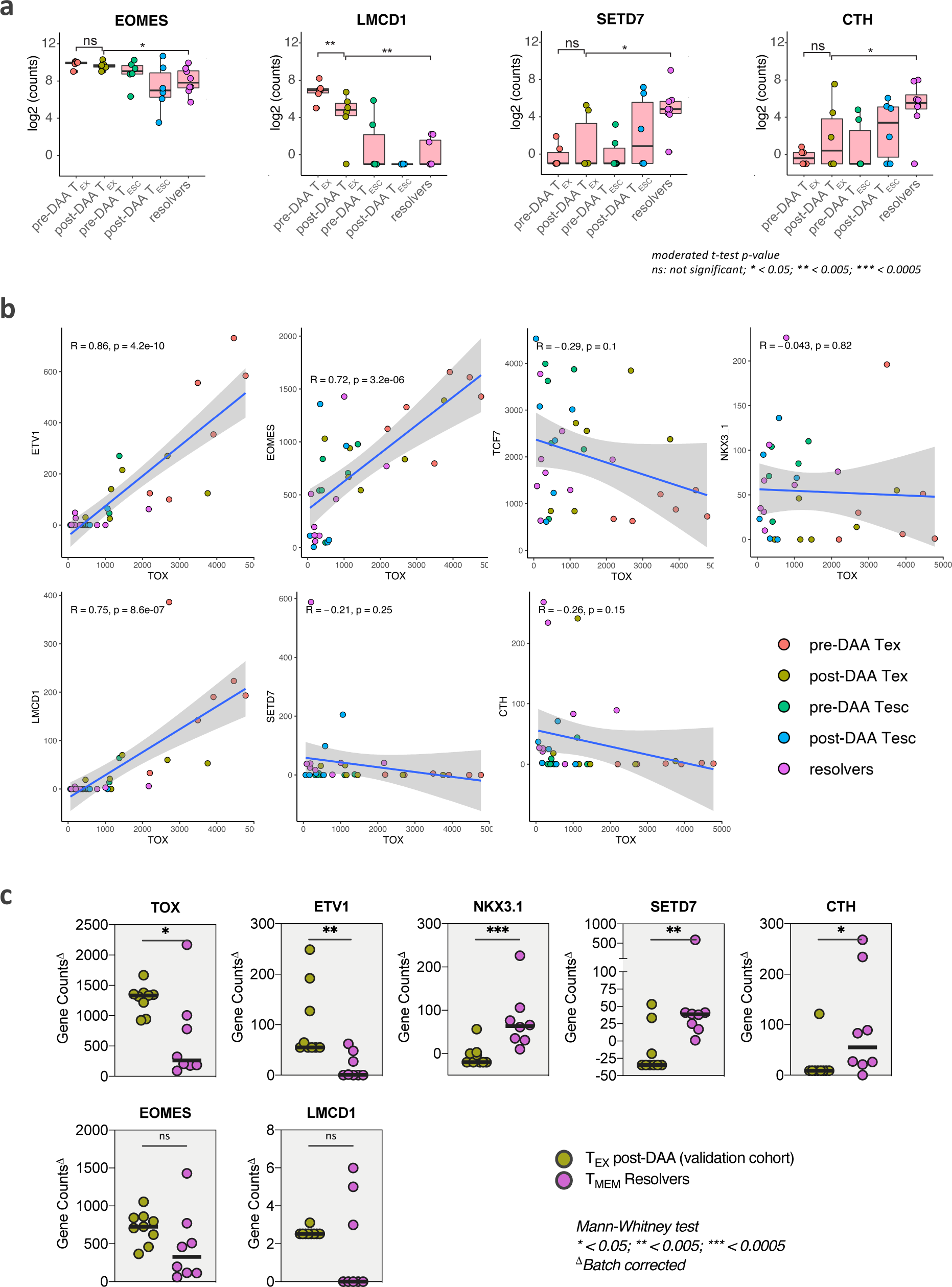
**a**, Expression (log_2_ counts) of EOMES, LMCD1, SETD7 and CTH in T_EX_ and T_ESC_ pre- and post-DAA (paired samples n=6) as well as in resolver T_MEM_ cells (n=8). **b**, Linear regression analysis to model the relationship in gene count expression of TOX and the other transcription factors and co-factors identified in Fig. 6d, by the different populations of HCV-specific CD8+ T cells, pre- and post-DAA therapy or after spontaneous resolution. Pearson correlation coefficient R and significance p (two-sided) values are reported from the linear regression analysis performed with R software. c, Gene count expression of TOX, ETV1, NKX3-1, SETD7, CTH, EOMES and LMCD1, by HCV-specific CD8+ T cells from a validation cohort of additional individual with T_EX_ post-DAA (n=9), as compared to resolver T_MEM_ cells (n=8), and following batch effect correction.

**Extended Data Table 1:** Patient demographic information and class I HLA types

| Patient ID | Sex | Age at Enrollment | Class I HLA type |  |  |  |  |  |
| --- | --- | --- | --- | --- | --- | --- | --- | --- |
|  |  |  | A1 | A2 | B1 | B2 | C1 | C2 |
| Chronically HCV-infected patients |  |  |  |  |  |  |  |  |
| 101 | Male | 29 | 0201 | 0301 | 1401 | 5101 | 0802 | 1402 |
| 102 | Male | 55 | 0201 | 0201 | 4402 | 5101 | 0401 | 0501 |
| 103 | Female | 56 | 0201 | 1101 | 1501 | 5201 | 0304 | 1202 |
| 104 | Male | 36 | 0201 | 2402 | 0801 | 4402 | 0501 | 0701 |
| 105 | Male | 28 | 0301 | 0301 | 0702 | 0702 | 0702 | 0702 |
| 106 | Male | 31 | 0101 | 0201 | 5101 | 5701 | 0602 | 1502 |
| 107 | Male | 27 | 0201 | 2501 | 1524 | 5701 | 0303 | 0602 |
| 108 | Female | 72 | 2601 | 3101 | 0702 | 4402 | 0501 | 0702 |
| 109 | Male | 42 | 2402 | 6601 | 0702 | 3801 | 0702 | 1203 |
| 110 | Male | 44 | 0201 | 0201 | 4402 | 4402 | 0501 | 0501 |
| 111 | Male | 39 | 0201 | 0301 | 0702 | 4001 | 0304 | 0702 |
| 112 | Male | 37 | 0201 | 2902 | 1402 | 3901 | 0802 | 1203 |
| 113 | Male | 24 | 0101 | 3101 | 0702 | 0702 | 0702 | 0702 |
| 114 | Male | 38 | 2402 | 3001 | 1302 | 4403 | 0602 | 1601 |
| 115 | Male | 42 | 0101 | 0201 | 1518 | 3502 | 0401 | 0704 |
| 118 | Female | 51 | 0101 | 0201 | 0801 | 1501 | 0303 | 0701 |
| 119 | Male | 47 | 0201 | 0301 | 0702 | 1402 | 0702 | 0802 |
| 120 | Male | 57 | 0101 | 2501 | 0702 | 0801 | 0701 | 0702 |
| 121 | Male | 32 | 0201 | 3004 | 0702 | 3502 | 0702 | 1402 |
| 123 | Female | 62 | 0301 | 1101 | 3501 | 4001 | 0304 | 0401 |
| HCV-spontaneous resolvers |  |  |  |  |  |  |  |  |
| BR-3000 | Male | 41 | 0201 | 1101 | 0702 | 4403 | 0702 | 1602 |
| 06-00K | Female | 22 | 0101 | 3201 | 2705 | 3503 | 0202 | 0401 |
| 09-038 | Female | 20 | 02:01 | 68:02 | 14:02 | 4402 | 0501 | 0802 |
| 12-055 | Male | 30 | 0201 | 6801 | 3508 | 5701 | 0401 | 0602 |
| 12-140 | Female | 37 | ND | ND | ND | ND | ND | ND |
| 09-033 | Female | 41 | 0101 | 0101 | 0801 | 4402 | 0501 | 0701 |
| BR-2350 | Male | 43 | 0101 | 1101 | 0801 | 3501 | 0401 | 0701 |
| 05-00Y | Female | 21 | 0201 | 2402 | 1501 | 4001 | 0303 | 0304 |
| 08-023 | Male | 27 | 0201 | 3001 | 2705 | 5201 | 0102 | 1202 |
| BR-320 | Female | 35 | 0201 | 3101 | 1501 | 5101 | 0102 | 1203 |

**Extended Data Table 2:** List of antibodies used for flow-cytometry analyses.

| Antibody and fluorochrome | Clone | Manufacturer | Reference |
| --- | --- | --- | --- |
| CD3-PE Dazzle 594 | UCHT1 | BIOLEGEND | 300450 |
| CD8-BUV-395 | RPA-T8 | BD BIOSCIENCES | 563795 |
| CD19-BUV-496 | SJ25C1 | BD BIOSCIENCES | 564655 |
| TCF-1-PE | S33-966 | BD BIOSCIENCES | 564217 |
| EOMES-FITC | WD1928 | EBIOSCIENCE | 11-4877-42 |
| T-bet-BV421 | O4-46 | BD BIOSCIENCES | 563318 |
| PD-1-BV421 | EH12.2H7 | BIOLEGEND | 329920 |
| CD95-BV421 | DX2 | BIOLEGEND | 305623 |
| TIGIT-PE | MBSA43 | EBIOSCIENCE | 12-9500-42 |
| 2B4-PerCP-Cy5.5 | C1.7 | BIOLEGEND | 329516 |
| BTLA-PE | MIH26 | BIOLEGEND | 344505 |
| CD160-AF488 | BY55 | EBIOSCIENCE | 53-1609-42 |
| KLRG1-BV605 | 2F1/KLRG1 | BIOLEGEND | 138419 |
| TIM-3-BV421 | F38-2E2 | BIOLEGEND | 345008 |
| CTLA-4-PE-Cy7 | L3D10 | BIOLEGEND | 349913 |
| LAG-3-PE | 3DS223H | EBIOSCIENCE | 12-2239-41 |
| CD27-APC-Cy7 | O323 | BIOLEGEND | 302815 |
| CD38-PE-Cy7 | HB7 | EBIOSCIENCE | 25-0388-42 |
| HLA-DR-BV605 | L243 | BIOLEGEND | 307640 |
| CD28-PE | CD28.2 | BIOLEGEND | 302908 |
| ICOS-APC-Cy7 | c398.4a | BIOLEGEND | 313529 |
| CD69-FITC | FN50 | BIOLEGEND | 310903 |
| CD71-AF700 | M-A172 | BD BIOSCIENCES | 563769 |
| CD137-AF700 | 4B4-1 | BIOLEGEND | 309816 |
| DNAM-1-FITC | DX11 | BD BIOSCIENCES | 559788 |
| CD161-BV510 | HP-3G10 | BIOLEGEND | 339921 |
| IB7-FITC | FIB504 | BIOLEGEND | 321212 |
| CD103-BV605 | Ber-ACT8 | BIOLEGEND | 350217 |
| CCR4-BV605 | L291H4 | BIOLEGEND | 359417 |
| CD62L-PerCP-Cy5.5 | DREG-56 | BIOLEGEND | 304824 |
| CD39-BV510 | A1 | BIOLEGEND | 328219 |
| CD26-PE-Cy7 | BA5b | BIOLEGEND | 302714 |
| CD73-PE-Cy7 | AD2 | BIOLEGEND | 344010 |
| Granzyme-B-BV510 | NGZB | BD BIOSCIENCES | 563388 |
| Perforin-PerCP-Cy5.5 | DG9 | BIOLEGEND | 308114 |
| KI67-AF700 | B56 | BD BIOSCIENCES | 561277 |
| CD57-PerCP-Cy5.5 | HNK-1 | BIOLEGEND | 359621 |
| CD127-BV421 | A019D5 | BIOLEGEND | 351309 |
| CCR7-APC-Cy7 | G043h7 | BIOLEGEND | 353211 |
| CD45RA-AF700 | HI100 | BIOLEGEND | 304120 |
| IFNγ-APC-Cy7 | 4S.B3 | BIOLEGEND | 502530 |
| TNFα-BV605 | MAb11 | BIOLEGEND | 502936 |
| IL-2-PE-Cy7 | MQ1-17H12 | BIOLEGEND | 500326 |
| CD107a-BV421 | H4A3 | BIOLEGEND | 328626 |

**Extended Data Table 3:** 1248 genes with significantly different expression levels were identified between the HCV-specific CD8 T cell populations from different disease states using limma package in R (adjusted P value cut-off = 0.05). Abbreviations used in the table are: R - resolvers; ConPre - preDAA exhausted T cells; ConPost - postDAA exhausted T cells; MutPre – preDAA escaped T cells; MutPost – postDAA escaped T cells. In addition, we have included data from the same individuals for the following non-specific CD8 T cell populations for comparison: CmPre – preDAA central memory T cells; CmPost – postDAA central memory T cells; CmRes - resolver central memory T cells.

